# KLF4 Recruits SWI/SNF to Increase Chromatin Accessibility and Reprogram the Endothelial Enhancer Landscape under Laminar Shear Stress

**DOI:** 10.1101/2020.07.10.195768

**Authors:** Jan-Renier A.J. Moonen, James Chappell, Minyi Shi, Tsutomu Shinohara, Dan Li, Maxwell R. Mumbach, Fan Zhang, Joseph Nasser, Daniel H. Mai, Shalina Taylor, Lingli Wang, Ross J. Metzger, Howard Y. Chang, Jesse M. Engreitz, Michael P. Snyder, Marlene Rabinovitch

**Author notes:** Co-first author (Jan-Renier A.J. Moonen, James Chappell).

## Abstract

Physiologic laminar shear stress (LSS) induces an endothelial gene expression profile that is vasculo-protective. In this report, we delineate how LSS mediates changes in the epigenetic landscape to promote this beneficial response. We show that under LSS, KLF4 interacts with the SWI/SNF nucleosome remodeling complex to increase accessibility at enhancer sites that promote expression of homeostatic endothelial genes. By combining molecular and computational approaches we discovered enhancers that loop to promoters of known and novel KLF4- and LSS-responsive genes that stabilize endothelial cells and suppress inflammation, such as *BMPR2* and *DUSP5*. By linking enhancers to genes that they regulate under physiologic LSS, our work establishes a foundation for interpreting how non-coding DNA variants in these regions might disrupt protective gene expression to influence vascular disease.

## Introduction

Regions of the arterial circulation exposed to uniform physiologic laminar shear stress (LSS) are associated with expression of vasculoprotective genes such as endothelial nitric oxide synthase (*eNOS*), whereas regions of low and disturbed shear stress, near arterial branches and bifurcations, are more prone to develop disease^1-5^. However, the nature and severity of vascular lesions depends on whether there are mitigating genetic and environmental factors that not only worsen the propensity to disease in vulnerable regions of the circulation, but also perturb the normal response to LSS.

Major effort has been directed at finding mechanosensing complexes in endothelial cells (EC) that respond to LSS to initiate a pattern of protective gene regulation that is altered in disease^6-8^. Krüppel-like factors (KLF) 2 and 4 have been identified as key transcription factors induced by LSS that control endothelial homeostatic gene regulation^9-13^. Beyond transcriptional activation at gene promotors, these factors might function as chromatin organizers to regulate genes by altering the chromatin landscape at distal enhancers. Knowing how the enhancer landscape changes in response to LSS provides a blueprint that will better inform how variants in non-coding DNA can influence vascular diseases, including pulmonary arterial hypertension (PAH), which is characterized by progressive occlusion of distal pulmonary arteries^14,15^. Pathogenic gene variants in *BMPR2* are the most common genetic risk factor for PAH, but have largely been associated with only 15% of patients with the familial form of the disease^16,17^, suggesting that variants in non-coding DNA regions might contribute to reduced BMPR2 levels in other patients with PAH.

Whereas variants in coding and proximal regulatory DNA regions can be readily related to dysfunction of a particular disease-causing gene, ascribing variants in distal non-coding regions to genes related to vascular disease has been more elusive. New bioinformatic approaches are overcoming this hurdle. For example, recently a variant in a distal enhancer that targets endothelin-1 was linked to five vascular diseases^18^. With more effort directed at whole genome sequencing in cohorts of patients such as those with PAH, establishing the functional significance of variants in non-coding DNA has become increasingly important. Moreover, variants identified in enhancers that are active under physiologic LSS might have the greatest adverse impact on gene regulation.

We therefore sought to map the enhancer landscape of pulmonary artery EC (PAEC) exposed to LSS by first relating changes in chromatin accessibility determined by ATAC-Seq, to gene expression assessed by RNA-Seq. We found an overall increase in chromatin accessibility, coinciding with the induction of protective genes and suppression of disease-related genes. Motif analyses of differentially accessible regions (DAR) revealed enrichment for KLF binding sites in regions with increased accessibility under LSS, which we confirmed by KLF4 ChIP-Seq. We demonstrated, using KLF gain-and-loss-of-function experiments, that KLF is required to mediate chromatin accessibility changes at proximal and distal sites, adding to the known role for KLF in transcriptional activation of flow responsive genes.

Consistent with this finding, we identified components of the SWItch/Sucrose Non-Fermentable (SWI/SNF) nucleosome remodeling complex that were recruited by KLF4 and required for KLF mediated gene regulation. H3K27ac HiC Chromatin Immunoprecipitation (HiChIP) ^19,20^ showed increased enhancer looping to LSS-responsive genes, often spanning multiple genes. Application of the Activity-by-Contact (ABC) algorithm^21^, which identifies regulatory elements of specific genes based upon chromatin accessibility and H3K27ac levels, predicted more than 70% of genes differentially expressed under LSS to be regulated by KLF4. These data provide a blueprint of the endothelial enhancer landscape under physiologic LSS, which informs future studies on how variants in non-coding DNA might impair protective gene expression and promote vascular disease.

## Results

### LSS alters chromatin accessibility, increasing expression of vasculo-protective genes

Genome-wide changes in chromatin accessibility were studied by ATAC-Seq in PAEC exposed to a physiologic level of 15 dyn/cm^2^ of LSS^5,22^ for 24h. Exposure to LSS increased accessibility of 4,699 genomic regions and 2,191 regions showed a loss of accessibility compared to PAEC cultured under static conditions (ST) (**Fig. 1a**). While a subset of these differentially accessible regions (DAR) contain gene promoters, most DAR are located in introns or at intergenic sites. (**Fig. 1b**). When assigning each DAR to its nearest gene, we found that the majority of DAR are located within 100 Kb of a transcription start site (**Fig. 1c**). To relate chromatin accessibility changes to altered gene expression, we performed RNA-Seq of PAEC cultured under LSS vs ST conditions. Induction of the LSS-responsive transcription factors KLF2 and KLF4 was confirmed by RNA-Seq and RT-qPCR (**Supplementary Fig. 1**). We observed 720 and 663 genes significantly up- or down-regulated, respectively, with LSS compared to ST conditions (**Fig. 1d).** Exposure to LSS resulted in induction of a quiescent vasculo-protective gene expression profile, as evident in enrichment for genes related to blood vessel development, improved barrier function and enhanced cell-matrix interaction (**Fig. 1e**, upper panel). In contrast, the gene expression profile of PAEC cultured under ST conditions is typical of a more activated and proliferative phenotype, including genes that promote cell migration, cell cycle molecules and genes related to oxidative stress (**Fig. 1e**, lower panel).

**Fig 1:**
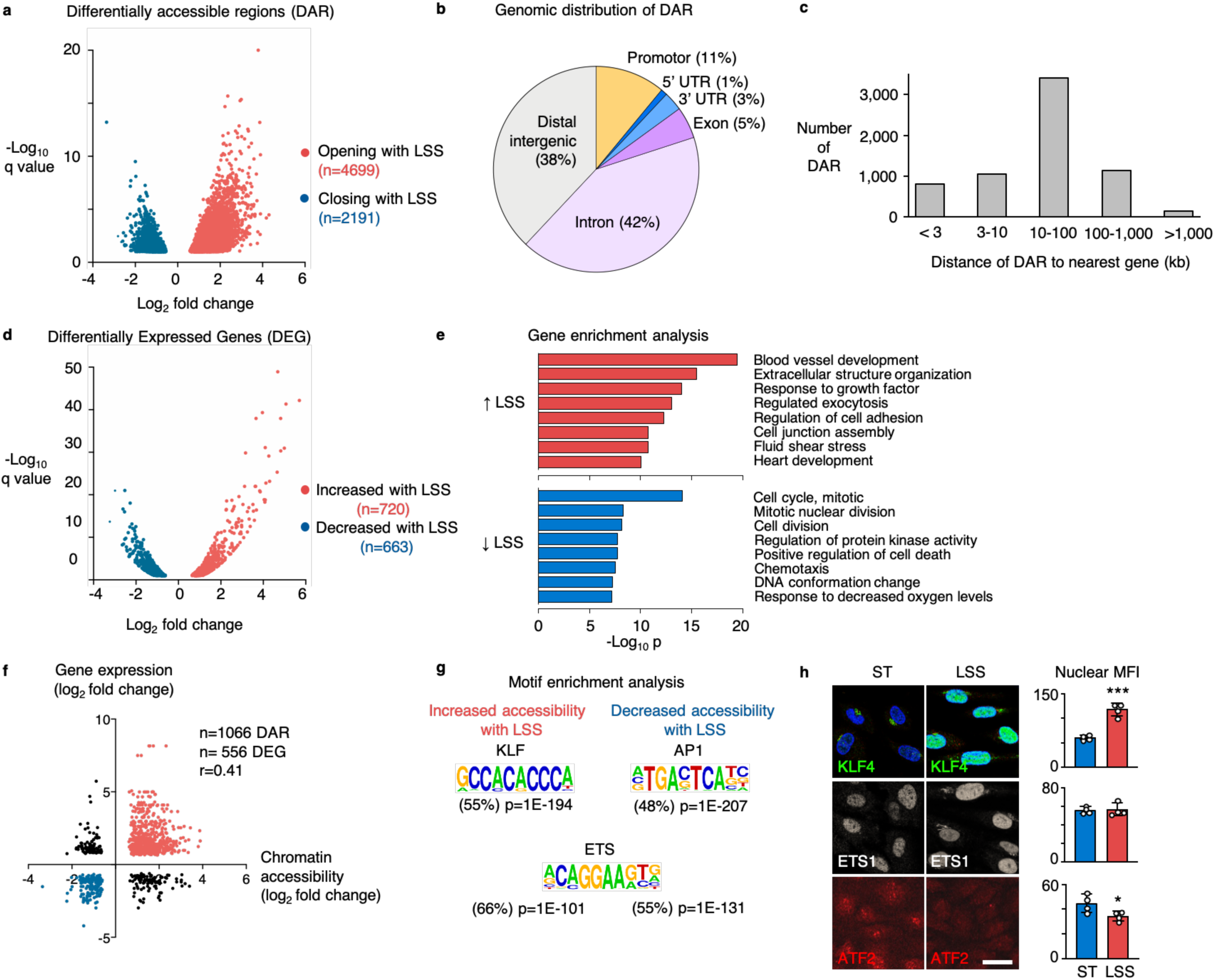
Laminar shear stress increases chromatin accessibility and vasculo-protective gene expression. **a**, Volcano plot showing the differentially accessible regions (DAR) determined by ATAC-Seq in PAEC that were exposed to 15 dyn/cm^2^ of LSS for 24h vs static controls. Each dot represents a DAR. Red dots are DAR with increased accessibility, blue dots are DAR with decreased accessibility under LSS. *n*=3 experimental replicates. *P* values were determined by the Wald test with Benjamini–Hochberg adjustment. **b**, Pie chart showing the annotation of DAR peaks, analyzed using Homer. **c**, Bar graphs showing the distance of DAR to its nearest gene, analyzed using Homer. **d**, Volcano plot showing the differentially expressed genes (DEG) determined by RNA-Seq in PAEC exposed to 15 dyn/cm^2^ of LSS for 24h vs Static controls. Each dot represents a DEG. Red dots are DEG with increased expression, blue dots are DEG with decreased expression under LSS. *n*=2 experimental replicates. *P* values were determined by the Wald test with Benjamini–Hochberg adjustment. **e**, Gene enrichment analysis using Metascape of genes that are induced by LSS (red bars) and genes that are repressed by LSS (blue bars). *P* values were determined by the hypergeometric test with Benjamin-Hochberg adjustment. **f**, Scatterplot showing a positive correlation of accessibility changes with the expression of its nearest differentially expressed gene. r=0.41 with *P*<0.0001 determined by Pearson R test. **g**, Motif enrichment of DAR with increased or decreased accessibility with LSS, analyzed using Homer. *P* values were determined by binomial test. **h**, Representative immunofluorescent images (left) with quantitation on the right, of PAEC that were exposed to 15 dyn/cm^2^ of LSS for 24h vs static controls (ST), showing the nuclear expression of KLF4 (green); ETS1 (grey) and ATF2 (red). Nuclei were stained with DAPI, (blue). *n*=4 experimental replicates. Bars indicate mean ± s.e.m. * *P*<0.05, *** *P*< 0.001 determined by Student’s two-tailed *t*-test. Scale bar, 20 μm.

When relating LSS-mediated DAR to their nearest gene, we found a positive correlation between chromatin accessibility changes and altered gene expression (r=0.41, *P*<0.0001) (**Fig. 1f**). Differentially expressed genes (DEG) associated with proximal accessibility changes were enriched for pathways related to extracellular matrix components, e.g., elastic fiber formation (*ELN, FBN1* and *FBLN1*), collagen formation (*COL3A1, COL4A1* and *COL5A1*), laminin interactions (*LAMA5, ITGB4* and *ITGB6*), and chondroitin sulfate/dermatan sulfate metabolism (*HYAL1, VCAN, GBE1, CHST3* and *CHST11*), as well as matrix metalloproteinases and disintegrins (*MMP15, MMP28, ADAM15, ADAMTS1* and *ADAMTS9*) and Insulin-like growth factor and Notch signaling components (*IGFBP5* and *IGFBP6*; and *HES1, JAG1* and *DLL1*).

DEG without DAR were enriched for pathways related to suppression of cell proliferation, e.g., a decrease in nucleosome assembly genes (*CENPA, CENPM, CENPN, CENPP* and multiple histone genes), and suppression of genes related to the resolution of sister chromatids (*PLK1* and *AURKB*), as well as several mini-chromosome maintenance (MCM) genes that are involved in the initiation of eukaryotic genome replication (*MCM2, MCM4, MCM5* and *MCM7*).

We then went on to identify the regulatory factors that mediate the chromatin accessibility changes by performing motif enrichment analyses on the DAR. In regions with increased accessibility under LSS, we found enrichment for motifs that are recognized by members of the KLF family (55% of DAR, p=1E-194), while activator protein-1 (AP1) motifs were most enriched in regions where accessibility was lost (48% of DAR, p=1E-207). Motifs for E26 transformation-specific (ETS) family members were present in about half of the DAR, in line with previous reports indicating that ETS is indispensable for endothelial gene regulation^23,24^ (**Fig. 1g**). We used immunofluorescence to study the nuclear expression of these transcription factor families under LSS and ST conditions, and confirmed the increased expression of KLF4 with LSS, while the AP1 family member activating transcription factor-2 (ATF2) was increased under static conditions. ETS1 was constitutively expressed under both conditions (**Fig. 1h**).

### KLF regulates chromatin accessibility of vasculoprotective genes under LSS

To determine whether KLF regulates changes in chromatin accessibility under LSS, we performed ATAC-Seq under conditions of KLF gain- or loss-of-function. For gain-of-function studies, we transduced PAEC with an adenoviral vector encoding a constitutively active mutant of mitogen-activated protein kinase kinase 5 (caMEK5), which leads to phosphoactivation of signal-regulated kinase 5 (ERK5) and the induction of KLF2 and KLF4^25,26^. PAEC transduced with caMEK5 showed a 14- and 22-fold increase in expression of KLF2 and KLF4, respectively, compared to controls transduced with a vector encoding green fluorescent protein (GFP) (**Supplementary Fig. 2**). Almost half of the DAR previously identified with LSS vs ST, were also DAR in caMEK5-transduced PAEC compared to GFP controls (n=3,211/6,890), and accessibility changes were highly correlated (r=0.89, *P*<0.0001) (**Fig. 2a**, left panel). When plotting the correlation of all LSS vs ST DAR with accessibility changes in caMEK5-transduced PAEC at those sites, we found a correlation of r=0.78 (P<0.0001) (**Supplementary Fig. 2**), indicating that activation of ERK5 and induction of KLF2/4 as previously described with LSS^26^ is largely responsible for the changes in chromatin accessibility. Similarly, the gene expression changes in caMEK5-transduced cells assessed by RNA-Seq strongly correlated with those observed with LSS (n=1,164, r=0.82, p<0,0001). (**Fig. 2a**, right panel).

**Fig 2:**
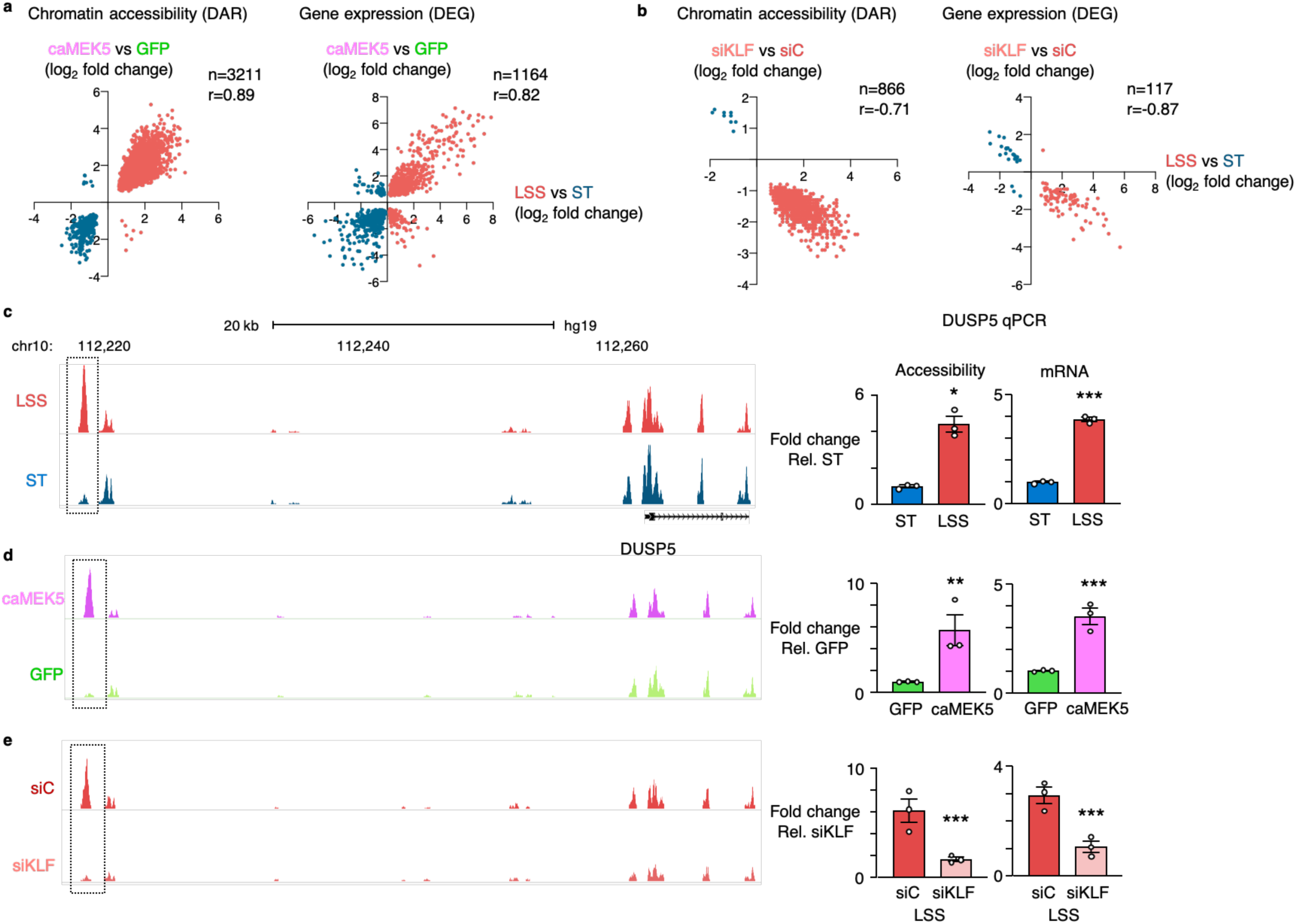
KLF regulates chromatin accessibility of vasculo-protective genes under LSS. **a**, PAEC were transduced with adenoviral vectors encoding caMEK5 or GFP as control. 90 h after transduction, chromatin accessibility changes were analyzed by ATAC-Seq and gene expression changes by RNA-Seq. Scatterplots show the correlation between changes in chromatin accessibility (left panel) and gene expression (right panel) in caMEK5 transduced PAEC compared to PAEC exposed to LSS. *n*=3 experimental replicates. Indicated values were calculated by Pearson R test, with *P*<0.0001. Each dot represents a DAR (left panel) or DEG (right panel). **b**, PAEC were treated with siRNA targeting KLF2 and KLF4 (siKLF), or non-targeting controls (SiC), 24h prior to exposing the cells to 15 dyn/cm^2^ of LSS vs ST conditions for an additional 24h. Chromatin accessibility changes were analyzed by ATAC-Seq and gene expression changes by RNA-Seq. Scatterplots show the correlation between changes in chromatin accessibility (left panel) and gene expression (right panel) in siKLF treated cells compared to untreated PAEC exposed to 15 dyn/cm^2^ of LSS for 24h. *n*=3 experimental replicates. Indicated values were calculated by Pearson R test, with *P*<0.0001. Each dot represents a DAR (left panel) or DEG (right panel). **c**, ATAC-Seq tracks showing a DAR upstream of *DUSP5* with increased accessibility under LSS vs ST control (left panel). Bar graphs showing the ATAC-qPCR analysis of the *DUSP5* DAR, and of *DUSP5* mRNA transcript levels by RT-qPCR (right panel). **d**, ATAC-Seq tracks showing increased accessibility of the DAR upstream of *DUSP5* in PAEC transduced with caMEK5 vs GFP controls. Bar graphs of the ATAC-qPCR and RT-qPCR analyses of caMEK5 transduced PAEC as described above (right panel). **e**, ATAC-Seq tracks of PAEC treated with siKLF as described above (left panel). Bar graphs of the ATAC-qPCR and RT-qPCR analyses of siKLF treated PAEC (right panel). For c, d and e, *n*=3 experimental replicates. qPCR data are shown as the mean ± s.e.m. * *P*<0.05, ** *P*<0.01, *** *P*< 0.001 by Student’s two-tailed *t*-test.

To further corroborate that KLF is required to alter chromatin accessibility under LSS, we performed loss-of-function experiments using RNAi targeting both KLF2 and KLF4 (siKLF). RNAi knockdown under LSS conditions reduced KLF2 and KLF4 expression levels by 79.7% and 69.8%, respectively (we note that this still corresponds to 4.9- and 7.8-fold higher expression compared to ST conditions, **Supplementary Fig. 2**). Upon RNAi knockdown, a subset of regions did not show the increase in accessibility that we previously observed with LSS (**Fig. 2b**, left panel) which coincided with a subset of genes not being induced (**Fig. 2b**, right panel). When plotting all LSS vs ST DAR with accessibility changes in siKLF-treated PAEC at those sites, we found a correlation of r=-0.73 (P<0.0001) (**Supplementary Fig. 2**).

We observed DAR near known endothelial flow responsive genes and KLF4 targets such as *eNOS* and thrombomodulin (*THBD*)^11^, but we also identified DAR associated with novel genes. An example is *DUSP5*, a dual specificity phosphatase that inhibits ERK1/2 signaling^27^. Via DUSP5 RNAi knockdown experiments under LSS, we found that DUSP5 is responsible for repressing multiple chemoattractant, antiviral and interferon genes (**Supplementary Fig. 2**). ATAC-Seq identified a potential regulatory DAR located 39 Kb upstream of the *DUSP5* transcription start site. ATAC-qPCR confirmed the increased accessibility at that site with LSS, which coincided with higher DUSP5 mRNA transcript levels (**Fig. 2c**). caMEK5 transduced cells had increased accessibility at the same regulatory region, which was confirmed by qPCR (**Fig. 2d**). When we performed RNAi to silence KLF2 and KLF4 prior to exposure to LSS, neither an increase in accessibility, nor the induction of *DUSP5* gene expression were observed (**Fig. 2e**).

### KLF4 interacts with the SWI/SNF nucleosome remodeling complex to increase chromatin accessibility

Having established that KLF regulates chromatin accessibility, we next sought to identify a chromatin remodeler with which it might interact to exert these changes. We transduced PAEC with an adenoviral vector encoding a Flag-tagged KLF4 mutant, or a GFP vector as control, after which affinity purification followed by mass spectrometry (AP-MS) was performed using anti-KLF and anti-Flag antibodies. We found, with both antibodies, that KLF4 co-purified with ten components of the SWI/SNF nucleosome remodeling complex, including the ATP-dependent helicase Brahma-Related Gene 1 (BRG1), also known as SMARCA4, and the invariable core subunit SMARCC2 (**Fig. 3a**). In addition to the SWI/SNF complex, we found KLF4 co-purified with several other chromatin remodelers that enhance chromatin accessibility and facilitate transcriptional activation, including the lysine demethylase KDM2A and Bromodomain Containing 4 (BRD4), as well as remodelers that decrease accessibility resulting in transcriptional repression, such as histone deacetylase HDAC2 and DNA methyltransferases DNMT1 and DNMT3A (**Fig. 3a**).

**Fig 3:**
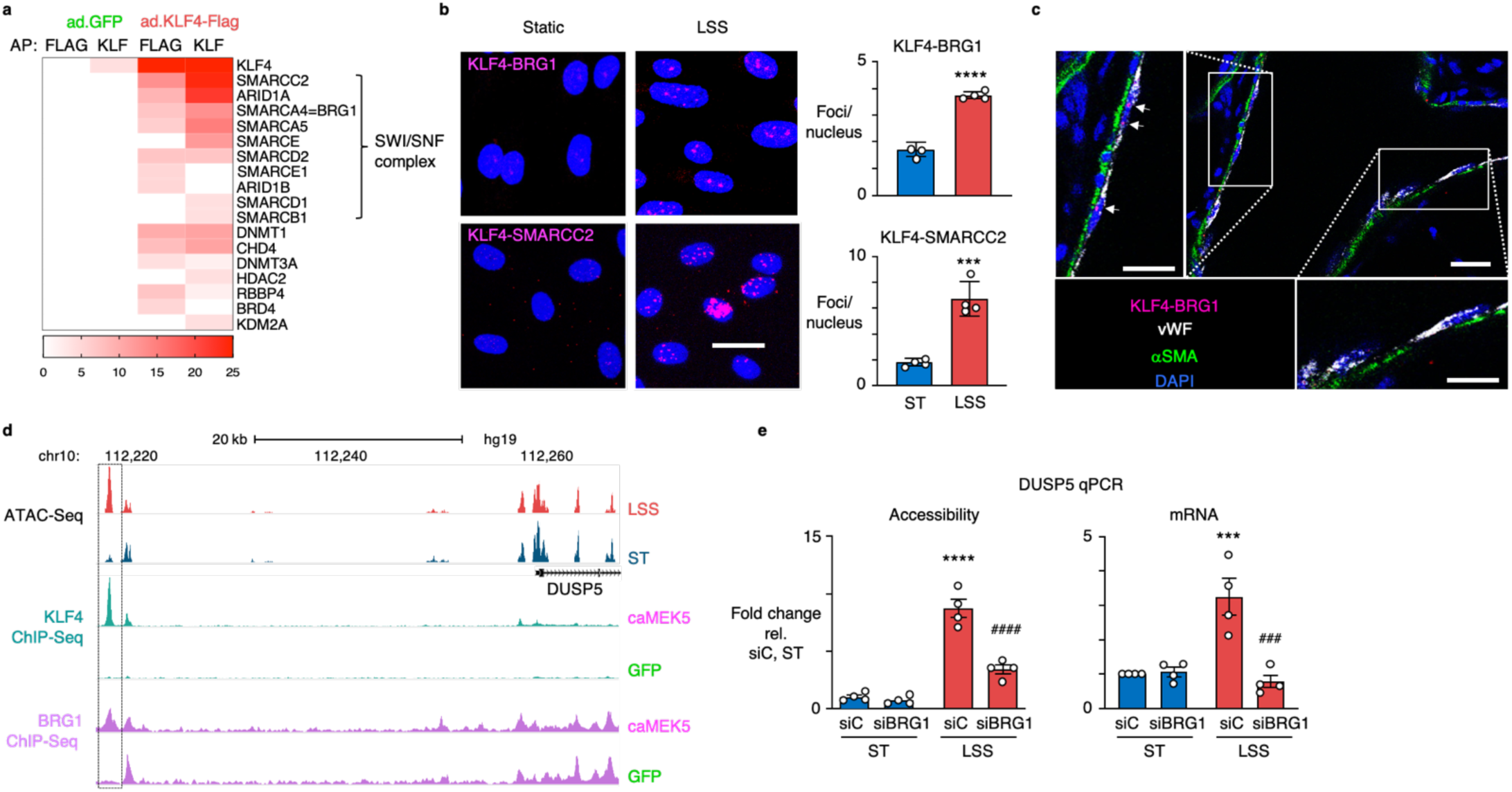
KLF4 interacts with the SWI-SNF nucleosome remodeling complex to increase chromatin accessibility. **a**, Heatmap showing the spectral counts obtained by Affinity Purification followed by Mass spectrometry (AP-MS) of PAEC transduced with adenoviral vectors encoding a Flag-tagged mutant of KLF4 or GFP control. AP was performed using anti-Flag antibodies (FLAG) and anti-KLF antibodies (KLF). Proteins were inferred from the peptides against the human UniProt using an FDR of 1%. *n*=2 experimental replicates. **b**, Representative images of Proximity Ligation Assays (PLA) (left panel) of PAEC exposed to 15 dyn/cm^2^ of LSS or ST conditions for 24h, show an interaction of KLF4 with BRG1 and SMARCC2 under LSS (magenta). Nuclei were stained with DAPI (blue). Number of foci per nucleus were quantified in 10 non-overlapping random fields of view per replicate (right panel). *n*=4 experimental replicates. Data shown as the mean ± s.e.m. *** *P*<0.001, **** *P*<0.0001 by Student’s two-tailed *t*-test. Scale bar, 20 μm **c**, Representative image of KLF4-BRG1 PLA in healthy rat lung tissue. The original PLA protocol was modified with longer incubation times to allow the reagents to fully penetrate the 350 μM thick sections. Note the interaction of KLF4 with BRG1 at straight sites of the vasculature that are exposed to LSS (upper left insert), while sites at bifurcations which are exposed to disturbed shear stress do not show the KLF4-BRG1 interaction (bottom right insert). KLF4-BRG1 interaction (magenta); vWF (grey, pseudo-colour); αSMA (green). Nuclei were stained with DAPI, (blue). Scale bar, 20 μm, and 10 μm in the higher magnifications. **d**, ATAC-Seq and KLF4 and BRG1 ChIP-Seq tracks showing enrichment for both factors at the DAR 39 Kb upstream of *DUSP5. n*=2 experimental replicates. **e**, PAEC were treated with siRNA targeting BRG1 (siBRG1) or with non-targeting controls (siC) prior to exposure to 15 dyn/cm^2^ of LSS or ST conditions for 24 h. Bar graphs indicate accessibility of the DAR 39 Kb upstream of *DUSP5*, and *DUSP5* gene expression, both assessed by qPCR. *n*=4 experimental replicates. Data are shown as the mean ± s.e.m. *** *P*<0.001, **** *P*<0.0001 siC LSS vs siC ST; ### *P*< 0.001, #### *P*<0.0001 siBRG1 LSS vs siC, by Student’s two-tailed *t*-test.

To confirm the interaction between KLF4 and the SWI//SNF complex in cells and tissue, we performed *in situ* proximity ligation assays (PLA) for KLF4 and either BRG1 or SMARCC2, which requires the epitopes of both proteins to be within 40 nm of each other to be detected. Indeed, in PAEC exposed to LSS, we observed the interaction between KLF4 and both SWI/SNF components (**Fig. 3b).** To determine if this interaction also occurs in the intact vessel wall, we modified the PLA protocol to allow us to study the KLF4-BRG1 interaction in the rat pulmonary vasculature. Using deep tissue imaging we focused on regions with bifurcations to allow us to compare straight parts of the vessel exposed to LSS, with branching points that are exposed to disturbed shear stress (**Supplementary Fig. 3**). The KLF4-BRG1 interaction was prominent at sites of LSS and much less apparent at regions with disturbed flow (**Fig. 3c**).

We then performed ChIP-Seq for KLF4 and BRG1 in caMEK5-versus GFP-transduced cells. At the DAR 39 Kb upstream of *DUSP5* that we had previously identified by ATAC-Seq, we found enrichment for both KLF4 and BRG1 (**Fig. 3d**). To confirm that BRG1 is required for the chromatin accessibility changes, we transfected PAEC with RNAi targeting BRG1 prior to exposure to LSS. Indeed, loss of BRG1 decreased chromatin accessibility of the DAR upstream of *DUSP5* as assessed by ATAC-qPCR, coinciding with reduced DUSP5 mRNA transcript levels (**Fig. 3e**).

### KLF4 and BRG1 co-occupied regions show a chromatin signature of enhancers

To study the genome wide chromatin accessibility changes co-regulated by KLF4 and BRG1, we intersected KLF4 and BRG1 peaks from the caMEK5 ChIP-Seq data with the ATAC-Seq data of PAEC exposed to LSS. Of the 8,404 regions that were enriched for both KLF4 and BRG1, 2,955 were found at DAR that open with LSS, representing 62.9% of all LSS DAR (**Fig. 4a**). In contrast, KLF4-BRG1 co-occupancy was only found at 4.2% of accessible regions that were not differentially accessible with LSS (**Supplementary Fig. 4**) and was only present in a small subset of DAR (2.1%) that lose accessibility under LSS (**Supplementary Fig. 4**). An increase in KLF4 binding at KLF4-BRG1 co-occupied DAR, correlated with increased accessibility (**Fig. 4b**).

**Fig 4:**
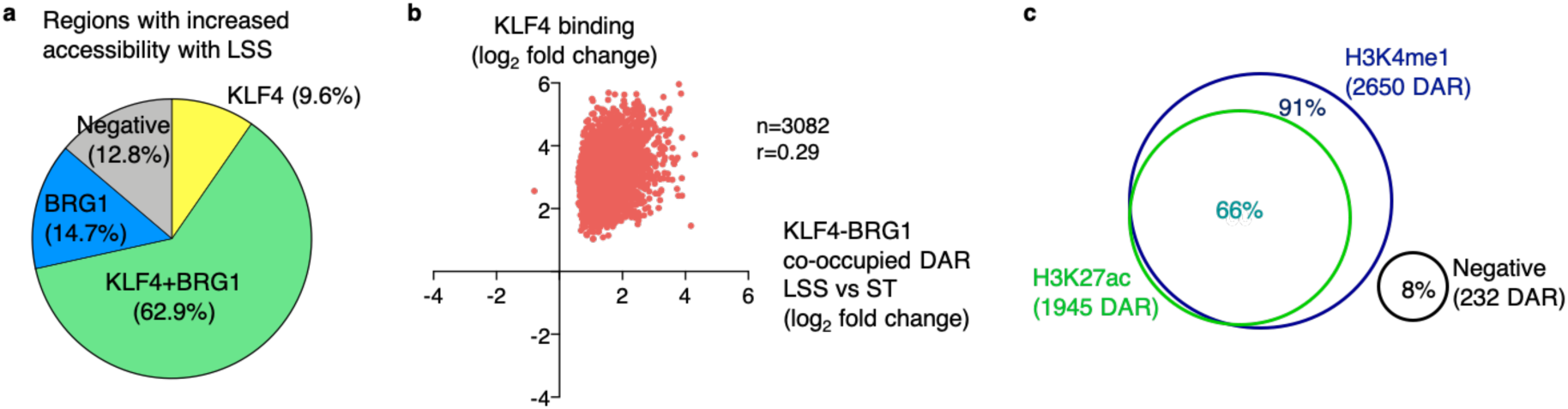
KLF4 and BRG1 co-occupy regions with increased accessibility under LSS, most of which are distal enhancers. **a**, Pie chart depicting the intersection of regions with increased accessibility under LSS by ATAC-Seq, with KLF4 and BRG1 ChIP-Seq data from PAEC transduced with caMEK5. Percentages indicate the fraction of DAR with increased accessibility under LSS vs ST that are differentially enriched for KLF4 and/or BRG1. **b**, Scatterplot showing the correlation between KLF4 binding and accessibility changes at LSS vs ST DAR co-occupied by KLF4-BRG1. Indicated values were calculated by Pearson R test, with *P*<0.0001. **c**, Venn diagram showing the percentage of KLF4-BRG1 co-occupied DAR with increased accessibility under LSS, that are enriched for H3K4me1 and H3K27ac. For a, b and c, *n*=2 experimental replicates.

Increased BRG1 peaks were found at 77.6% of DAR that open with LSS (**Fig. 4a**), as well as at 53.7% of DAR that open under ST conditions (**Supplementary Fig. 4**), but only in 13.5% of regions that were not differentially accessible (**Supplementary Fig. 4**), demonstrating that BRG1 binding coincides with increased chromatin accessibility even under static conditions. Confirmatively, when plotting changes in BRG1 binding at LSS vs ST DAR, we found increased BRG1 binding correlates with an increase in accessibility, and a decrease in BRG1 with loss of accessibility (r=0.89, p<0.0001) (**Supplementary Fig. 4**). BRG1 levels were twice as high in regions that were co-occupied by KLF4 compared to the average peak intensity of BRG1 across the genome, suggesting that KLF4 is recruiting BRG1 to those sites (**Supplementary Fig. 4)**.

To further characterize KLF4-BRG1 co-occupied regions, we performed ChIP-Seq for the H3K4me1 enhancer mark, as well as for H3K27ac that is typically present at active enhancers and promotors. The majority of KLF4-BRG1 co-occupied regions were enhancers marked by H3K4me1 (91%, **Fig. 4c**), and two thirds of those were active, as marked by H3K27ac (66%, **Fig. 4c**).

### KLF4 regulates gene expression by binding at enhancer loops

Thus far, we have related chromatin accessibility changes to gene expression based on the nearest gene. However, we found a direct relationship in only a subset of DEG, likely because several enhancers target a particular gene, and also because enhancer loops can skip promotors of proximal genes to target a more distal gene. We therefore applied two complementary strategies to map enhancer-promotor regulation: H3K27ac HiC Chromatin Immunoprecipitation (HiChIP), which measures the frequencies of 3D contacts between enhancers and promotors^19,20^, and the Activity-By-Contact (ABC) algorithm, that predicts enhancer-promotor interactions based on ATAC-Seq and H3K27ac ChIP-Seq data^21^.

Significant chromatin loops identified by HiChIP were determined using FitHiChIP^28^. At a one Kb resolution, we identified 4,698 enhancer-promotor (EP) loops, of which 1,299 were differential between LSS and ST conditions. We then integrated these data with KLF4 ChIP-Seq to study KLF4 binding at enhancer anchors, and found differential KLF4 binding at 856 enhancer anchors that target 325 DEG (**Fig. 5a**). An example is an enhancer loop that spans several genes to target the promoter of *BMPR2*. Mutations in the coding and proximal regulatory regions of this gene have been identified in patients with PAH^16,17^, but there might also be mutations in this distal enhancer that have not previously been linked to *BMPR2.* We identified a KLF4 bound EP loop with increased H3K27ac levels at the enhancer anchor located 1.3 Mb downstream of the *BMPR2* transcription start site (**Fig. 5b**, left panel), that is associated with a 2-fold increase in *BMPR2* mRNA levels under LSS (**Fig. 5b**, right panel). KLF4 also occupied enhancer anchors of loops that were lost with LSS, associated with a reduction in H3K27ac levels at those sites and a decrease in target gene expression. These include enhancer anchors of two EP loops that we identified to target *endothelin-1 (EDN1)*, a potent vasoconstrictor that is elevated in patients with PAH, and a major therapeutic target for the disease^29,30^. In caMEK5-transduced cells, there was increased binding of KLF4 at these sites with a concomitant decrease in H3K27ac levels (**Fig. 5c**, left panel). Based on our AP-MS data (**Fig. 3a**), we speculate that under this condition, KLF4 might interact with HDAC2 to inactivate the enhancer, resulting in decreased *EDN1* mRNA levels (**Fig. 5c**, right panel). In smooth muscle cells, the interaction between KLF4 and HDAC2 promotes transcriptional silencing of SM22α^31^.

**Fig 5:**
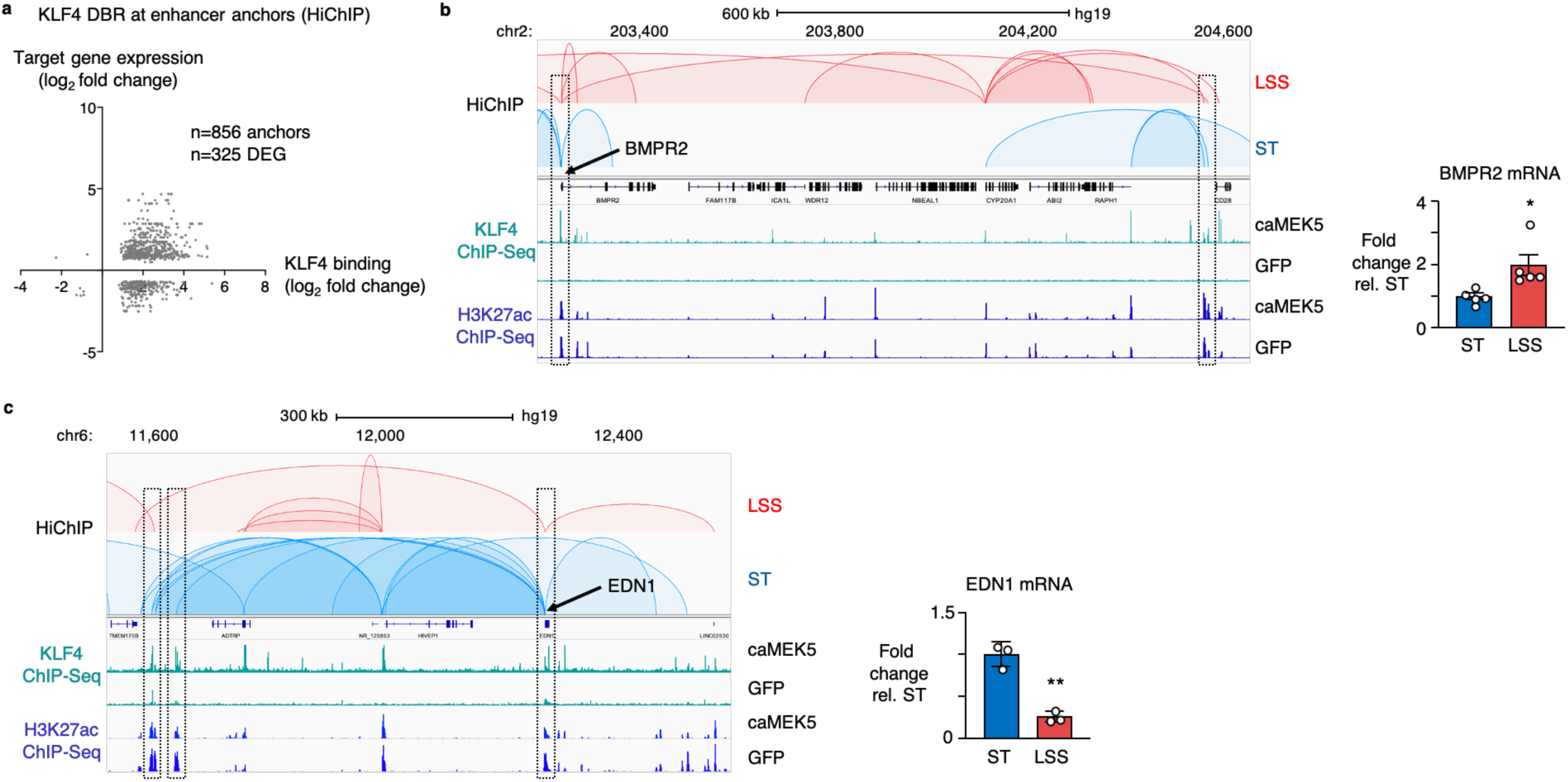
KLF4 modulates gene expression through changes in the enhancer landscape identified by H3K27ac HiChIP. **a**, Scatterplot showing the relation between differential KLF4 binding at enhancer anchors identified by HiChIP, with target gene expression. KLF4 ChIP-Seq was performed with PAEC transduced with caMEK5 vs GFP controls. HiChIP enhancer anchors were derived from PAEC exposed to 15 dyn/cm^2^ of LSS vs ST conditions for 24 h. *n*=2 experimental replicates for KLF4 ChIP-Seq; *n*=3 experimental replicates for HiChIP. **b**, HiChIP and ChIP-Seq tracks illustrating an example of a KLF4 occupied distal enhancer loop, spanning 1.3 Mb, that targets *BMPR2* (left panel). *BMPR2* mRNA was determined by RT-qPCR in PAEC exposed to 15 dyn/cm^2^ of LSS vs ST conditions for 24 h, and shown normalized to ST expression levels (right panel). *n*=5 experimental replicates. Data are shown as the mean ± s.e.m. * *P*<0.05 by Student’s two-tailed *t*-test. **c**, HiChIP and ChIP-Seq tracks showing loss of two distal enhancer loops, spanning 680 Kb and 638 Kb, that target *EDN1* and are occupied by KLF4 in PAEC transduced with caMEK5, coinciding with loss of H3K27ac (left panel*). EDN1* mRNA was determined by RT-qPCR in PAEC exposed to 15 dyn/cm^2^ of LSS vs ST conditions for 24 h and shown normalized to ST expression levels (right panel). *n*=3 experimental replicates. Data are shown as the mean ± s.e.m. ** *P*<0.01 by Student’s two-tailed *t*-test.

The ABC algorithm predicts which distal elements regulate which genes, considering elements located at both shorter (<10 Kb) and longer (>10 Kb) ranges^21^. The algorithm scores elements based on chromatin accessibility determined by ATAC-Seq, and the strength of the H3K27ac enhancer signal at those sites, and has been validated by perturbing thousands of putative enhancers using CRISPR interference^21^. The algorithm predicted 996/1,383 DEG under LSS to be regulated by enhancer elements that were occupied by KLF4 (**Fig. 6a**). ABC confirmed most of the enhancer loops that were identified by HiChIP, for example, those targeting *EDN1* described above, as well as one targeting the *SMAD5* promotor (**Fig. 6b**), and also identified enhancer loops that were not discovered by HiChIP, such as an enhancer for the Notch target gene *HES2* (**Fig. 6c**).

**Fig 6:**
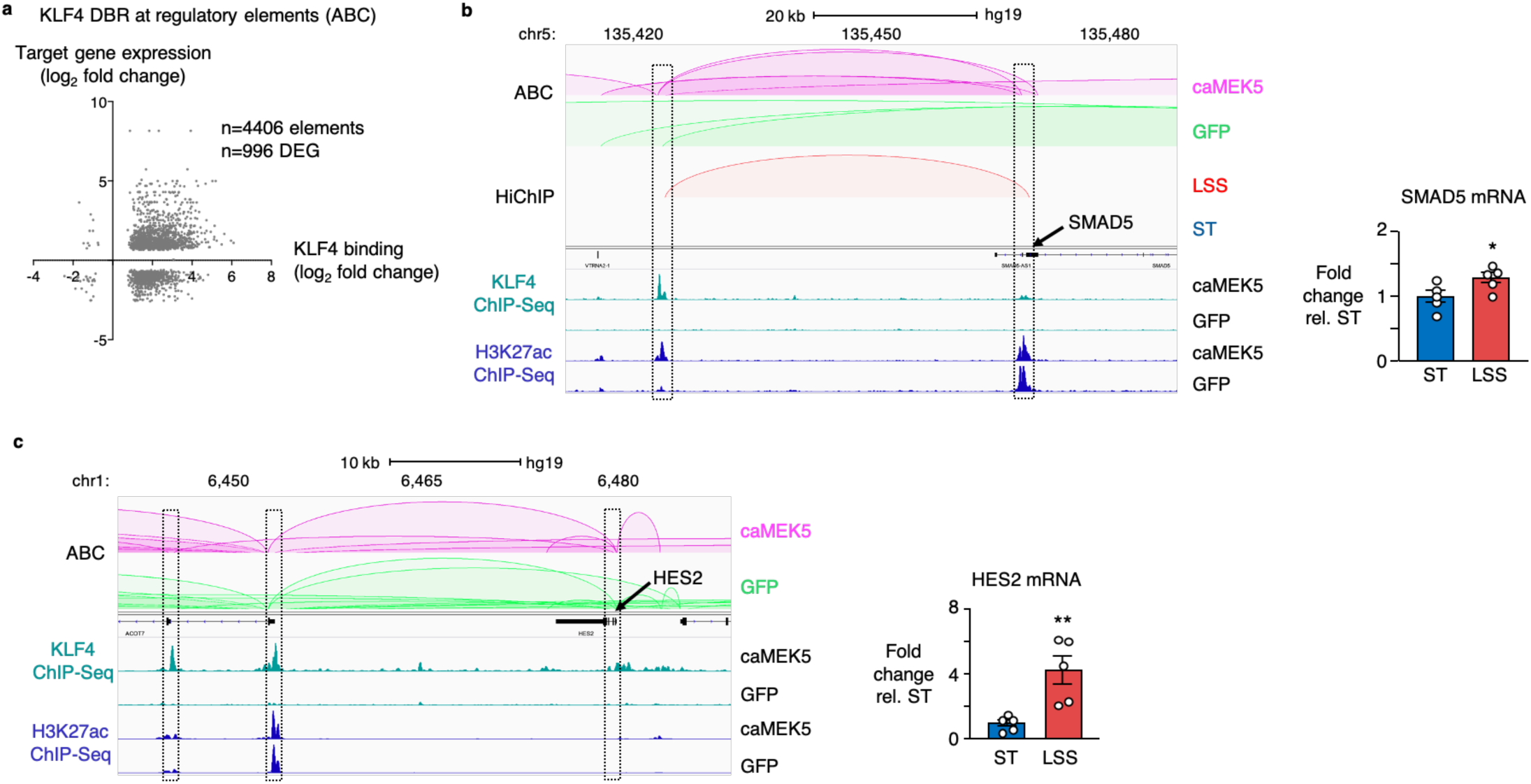
The ABC algorithm predicts that most LSS-responsive genes are regulated by KLF4 binding at enhancers. **a**, Scatterplot showing the relation between differential KLF4 binding at enhancer anchors identified by ABC, with target gene expression. KLF4 ChIP-Seq was performed with PAEC transduced with caMEK5 vs GFP controls. For ABC, H3K27ac CHIP-Seq and ATAC-Seq analyses from PAEC transduced with caMEK5 vs GFP were used as input. *n*=2 experimental replicates for all. **b**, ABC, HiChIP and ChIP-Seq tracks illustrating a KLF4 bound enhancer loop that spans 45 Kb and targets *SMAD5* that was identified by both HiChIP and ABC (left panel). *SMAD5* mRNA was determined by RT-qPCR in PAEC exposed to 15 dyn/cm^2^ of LSS vs ST conditions for 24 h, and shown normalized to ST expression levels (right panel). *n*=5 experimental replicates. Data are shown as the mean ± s.e.m.* *P*<0.05 by Student’s two-tailed *t*-test. **c**, ABC and ChIP-Seq tracks showing two KLF4-occupied proximal enhancer loops 27 Kb and 34 Kb upstream of the *HES2* target gene (left panel). *HES2* mRNA was determined by RT-qPCR in PAEC exposed to 15 dyn/cm^2^ of LSS vs ST conditions for 24 h, and shown normalized to ST expression levels (right panel). *n*=5 experimental replicates. Data are shown as the mean ± s.e.m. ** *P*<0.01 by Student’s two-tailed *t*-test.

## Discussion

KLF2 and KLF4 are well-recognized LSS-induced transcription factors that control endothelial homeostatic gene regulation^9-13^. Our studies indicate that KLF4, beyond its previously established role as a transcriptional activator of vasculoprotective gene expression, acts as a chromatin organizer by recruiting the SWI/SNF chromatin remodeling complex to alter chromatin accessibility and regulate the endothelial enhancer landscape.

Shear forces exerted by blood flow are sensed by endothelial cells and converted to protective or vulnerable gene expression profiles through mechanotransduction. Identification of mechanosensing complexes and their downstream signaling pathways has been the subject of extensive investigation and has greatly improved our understanding of how spatiotemporal changes in hemodynamics affect endothelial function and vascular homeostasis^6-8^. More recently, the epigenetic regulation of shear stress-mediated gene expression has gained increasing attention. Illi et al. were the first to report the importance of H3 and H4 histone modifications in the regulation of LSS-induced gene expression^32^. Many studies followed that investigated the role of histone acetyltransferases and deacetylases, as well as the histone methyltransferase EZH2, in mediating shear stress induced gene expression changes both *in vitro* and *in vivo*^33-37^. Other studies have addressed the role of DNA methylation and the contribution of DNA methyltransferases (DNMT) in the regulation of gene expression by shear stress^38-40^, including the identification of DNMT3A-dependent KLF4 promotor hypermethylation by disturbed flow^41^.

He and colleagues recently demonstrated that KLF4 regulates Inositol 1,4,5-trisphosphate receptor type 3 (IPTR3) expression by increasing chromatin accessibility and H3K27 acetylation at the ITPR3 promotor region^42^. In addition, many microRNAs are shear stress-regulated^43^, perhaps most notably, miR-92a, that targets KLF2 and KLF4 under conditions of disturbed flow^44,45^. More recently, long non-coding RNAs (lncRNAs) have been studied as epigenetic regulators of shear stress-induced gene expression, leading to the discovery of spliced-transcript endothelial-enriched lncRNA (STEEL)^46^ and a lncRNA that enhances eNOS expression (LEENE)^47^.

Yet despite these advances, no studies have comprehensively mapped the endothelial regulatory landscape under physiologic LSS. While transcription factor binding to gene promotors is sufficient to drive basal levels of transcription^48^, activation of enhancers is required for full transcriptional activity^49^, and to ensure phenotypic robustness^50^. In cultured cells, persistent changes in the epigenome may be particularly revealing of mechanisms that converge to cause disease^51^.

To investigate LSS-induced changes to the chromatin landscape, we performed ATAC-Seq on PAEC that were exposed to physiologic LSS of 15 dyn/cm^2^ for 24h. Only a small subset of DAR was located in gene promotors, the majority being putative regulatory elements located in introns or intergenic sites. ChIP-Seq analyses of H3K4me1 and H3K27ac confirmed that the majority of these DAR have a chromatin signature typical of enhancers. Motif enrichment analysis of DAR with increased accessibility with LSS pinpointed the KLF transcription factor family as potential regulators of the accessibility changes.

KLF4, in particular, has gained a lot of interest, being one of the four original reprogramming factors used by Takahashi and Yamanaka to create induced pluripotent stem cells (iPSC)^52^. This finding instigated further investigations into its role as a pioneer factor. Upon initial engagement of closed chromatin, pioneer factors recruit chromatin remodelers to allow stabilization of accessible chromatin, enabling other transcription factors to bind alongside them and recruit transcriptional machinery^53^. *In vitro*, purified KLF4 binds nucleosomes, and *in vivo* KLF4 preferentially targets closed chromatin enriched for condensed nucleosomes^54^. Our KLF4 ChIP-Seq data corroborate these findings, showing KLF4 binding at 72.5% of regions that have increased accessibility versus 19.2% of regions that are constitutively accessible.

In the induction of pluripotency, KLF4 predominantly binds distal enhancers of genes that promote reprogramming, unless it partners with c-myc in which case it tends to bind promotors^55^. This was confirmed in our data set, in which 70.3% of KLF4 binding occurs outside of gene promotors that are within 3 Kb of transcription start sites (data not shown). It would be interesting to identify which transcriptional co-factors guide KLF4 to either distal enhancers or gene promotors. Given the enrichment for the ETS motifs in DAR, and the well-established roles for ETS factors in regulation of endothelial gene expression^23,24^, we speculate that distinct ETS family members may play important roles in guiding these processes.

Besides DNA methylation and histone modifications, ATP dependent chromatin remodeling processes such as the mammalian SWI/SNF complex, are important regulators of chromatin accessibility by disrupting histone–DNA contacts^56^. The SWI/SNF remodeling complex can assist transcriptional activators to access closed chromatin^57-59^ or facilitate subsequent transcriptional activation^60^. For example, in mouse embryonic stem cell differentiation, the pioneer factor Forkhead Box A2 (FOXA2) binds regions of closed chromatin and recruits SWI/SNF to regulate nucleosome depletion and to promote endodermal differentiation^61^. More recently, BRG1, one of two mutually exclusive ATPases that functions as the catalytic subunit of the SWI-SNF complex^62^, was related to reprogramming of iPSCs by increasing accessibility of pluripotency enhancers^63^. Alver and colleagues described SWI/SNF as required for maintenance of lineage specific enhancers^64^. While BRG1 co-localizes with pluripotency factors OCT4, SOX2 and NANOG in embryonic stem cells^65^, to our knowledge there have been no prior reports of KLF4 and SWI/SNF co-localization or interaction, other than a high throughput lentiviral proteomics study that identified an interaction between KLF4 and Brahma (BRM) which is closely related to BRG1, but mutually exclusive^66^. Because BRG1 is not regulated by shear stress, we speculate that under conditions of disturbed shear stress BRG1 might interact with AP1 family members, as we found enrichment for these motifs in DAR that are losing accessibility with LSS. Indeed, our preliminary PLA studies found AP1 member ATF2, which is increased in endothelial cells in atherosclerotic lesions^67^, to interact with BRG1 and SMARCC2 under ST conditions but not under LSS (data not shown).

Since physical distance is an important denominator in the likeliness that a regulatory element targets a certain gene, we initially related DAR to their nearest gene. While accessibility changes generally correlated with gene expression changes, this could only explain a subset of gene regulation. The seminal work by Dekker and colleagues, who first described chromosome confirmation capture (3C)^68^, instigated further development of 3C-derived methods, including landmark studies applying Hi-C^69,70^. This greatly improved our understanding of the 3D genome and helped explain how regulatory elements that are megabases away from their target promotors in the linear genome, are in fact in close physical proximity. To map the endothelial enhancer landscape and study the contribution of KLF4 and BRG1 in regulating enhancer activity and target gene expression, we applied two distinct but complementary strategies to study enhancer-promotor (EP) looping, i.e., H3K27ac HiChIP^19,20^ and the Activity-by-Contact model (ABC)^21^. The experimental H3K27ac HiChIP approach detects the frequency of 3D contacts and identified many long-range EP loops that often span genomic regions containing multiple genes. The computational ABC algorithm, that was extensively validated by thousands of CRISPRi perturbations^21^, confirmed most of the EP loops that were identified by HiChIP and discovered many additional EP loops. ABC predicted more than 70% of differentially expressed genes to be regulated by KLF4-bound enhancers, reinforcing the pivotal role of KLF4 as the mediator of the vasculoprotective effect of LSS. The importance of KLF4 in regulating the endothelial enhancer landscape is further supported by a recent study that uncovered a critical role for KLF4 in organizing the pluripotency-associated enhancer network^71^.

In most cases, KLF4 binding at enhancers resulted in increased target gene expression. However, in some, such as the EP loops targeting endothelin-1, KLF4 binding resulted in reduced H3K27ac and target gene expression, possibly by recruiting histone deacetylases such as HDAC2 that we identified to interact with KLF4 by AP-MS, and that others have shown to interact with KLF4 in smooth muscle cells^31^. These opposing changes in gene regulation reflect the diversity of remodelers that we identified as KLF4 interactors and warrant further study to understand how KLF4 binding results in different outcomes at different sites.

Our findings can be leveraged to better relate genetic variants in non-coding regions identified by genome-wide association studies (GWAS) to protective or pathogenic gene expression. Most single nucleotide polymorphisms (SNPs) identified by GWAS occur in non-coding regions^72^, and individually often contribute small or indirect effects to complex traits. Knowing the endothelial enhancer landscape under physiologic LSS, will facilitate linking variants identified by GWAS to specific genes, and to uncover novel genes related to cardiovascular disease, especially in rare conditions such as PAH.

## Contributions

J-R.M., J.C., M.P.S. and M.R. conceived the project and designed the research studies. J-R.M. and J.C. performed the experiments and analyzed the data. M.S. and D.L. provided technical assistance with the ChIP-Seq studies. T.S. performed RT-qPCR analyses. M.R.M. and H.Y.C. assisted with the HiChIP studies. F.Z. and R.J.M. provided assistance with the tissue imaging studies. J.N. and J.M.E. assisted with the ABC algorithm. D.H.M. assisted with the in vitro PLA studies. S.T. provided assistance with cell cultures. L.W. assisted with preparing tissues for imaging. J-R.M. and M.R. prepared the manuscript.

## Acknowledgements

We thank Drs. Caiyun Grace Li and Aiqin Cao (Stanford University) for their advice related to the AP-MS studies; Ms. Patricia del Rosario (Stanford University) for providing clinical information related to the cell and tissue samples, and Dr. Tushar Desai (Stanford University) for access to the Leica confocal microscope. We greatly appreciate the editorial assistance of Dr. Michal Bental Roof, and the administrative help of Ms. Michelle Fox. We are indebted to the Pulmonary Hypertension Breakthrough Initiative (PHBI), as the source of cells from unused donor lungs. The PHBI is funded by NIH/NHLBI R24 HL123767 and the Cardiovascular Medical Research and Education Fund (CMREF) UL1RR024986. De-identified demographic and clinical data were supplied by the Data Coordinating Center at the University of Michigan. Sequencing was performed by the Genomics and Personalized Medicine Sequencing Center, supported by award number NIH S10OD025212, and NIH/NIDDK P30DK116074. Mass spectrometry was performed at the Vincent Coates Foundation Mass Spectrometry Laboratory, Stanford University Mass Spectrometry, supported by award number S10RR027425 from the National Center for Research Resources. This work was supported by NIH/NHLBI R01 HL122887 (Drs. Marlene Rabinovitch and Michael Snyder) and R01 HL074186 (Dr. Marlene Rabinovitch). Dr. Jan-Renier Moonen was supported by funds from the California Tobacco-Related Disease Research Program of the University of California, award number 27FT-0039, and the Netherlands Heart Foundation, award number 2013T116. Dr. Marlene Rabinovitch is additionally supported by the Dwight and Vera Dunlevie Chair in Pediatric Cardiology at Stanford University.

## Methods

### Cell culture studies

Primary human pulmonary artery endothelial cells (PAEC) were either commercially obtained (PromoCell) or harvested from unused donor control lungs obtained though the Pulmonary Hypertension Breakthrough Initiative (PHBI) funded by NIH (R24 HL123767) and the Cardiovascular Medical Research and Education Fund (CMREF; UL 1RR024986). De-identified demographic and clinical data were obtained from the Data Coordinating Center at the University of Michigan (**Supplementary Table 1**). PAEC were grown in commercial EC media containing 5% FBS (Sciencell) in a 5% CO2 air atmosphere and used at passages 3–7. Cells were routinely tested for mycoplasma contamination. For shear stress experiments, PAEC were seeded in flow chamber slides (μ-Slide I 0.4 mm ibiTreat; Ibidi) and grown to confluence before exposure to 15 dyn/cm^2^ of unidirectional uniform laminar shear stress (LSS) for 24 h. LSS was generated using the Ibidi Perfusion System (Ibidi). Static controls were performed simultaneously with the shear stress experiments, cultured on standard tissue culture treated plates or dishes using the same EC media (Sciencell).

### RNAi

PAEC were transfected with ON-TARGETplus SMARTpool siRNA targeting KLF2 (L-006928-00-0005, Dharmacon), KLF4 (L-005089-00-0005, Dharmacon) and DUSP5 (L-003566-00-0005, Dharmacon) or ON-TARGETplus non-targeting pool (D-001810-10-05, Dharmacon) as siControl. Transfection was performed using Lipofectamine RNAiMAX in Opti-MEM 1 reduced serum medium (ThermoFisher) for 7 h, after which media were changed to regular ECM (Sciencell). 48 h after the start of transfection transfected PAEC were exposed to LSS or ST conditions for 24 h.

### Plasmids

PAEC were transduced with adenoviral constructs encoding a constitutive active mutant of dual specificity mitogen-activated protein kinase 5 (caMEK5) (#000101A, Applied Biological Materials Inc); Flag-tagged KLF4 (#VH829440, Vigene Biosciences) or GFP control (AVP004, GenTarget Inc) for 12 h after which cells were allowed to recover for 90 h before being harvested for subsequent experiments.

### ATAC-seq sample preparation and data analysis

ATAC-seq was performed as described in Buenrostro et al., 2015^73^. Briefly, endothelial cells were trypsinized to create a single-cell suspension. After counting, nuclei were isolated from 100,000 cells and sequencing adapters were transposed for 30 minutes at 37°C using 5 μl of TDE1 (Nextera Tn5 transposase, Illumina). After PCR and gel purification, libraries were subjected to 2×151 paired end sequencing on the Illumina HiSeq 4000 to obtain an average of approximately 58 million uniquely mapped reads per sample (Stanford Center for Genomics and Personalized Medicine, supported by NIH grant S10OD020141). The resulting data were processed using the Kundaje Lab ATAC-seq processing pipeline (https://github.com/kundajelab/atac_dnase_pipelines). Briefly, this pipeline takes FASTQ files as input, and outputs peak calls (accessible regions, AR). Alignments to AR were counted using DiffBind v2.4.8 (https://rdrr.io/bioc/DiffBind/) to produce a count matrix. Differentially accessible regions (DAR) were detected using DESeq2 (https://bioconductor.org/packages/release/bioc/html/DESeq2.html)^74^ with a P-value cutoff of <0.1. The HOMER (http://homer.salk.edu/homer/) function findMotifsGenome was used with default parameters to search for motif enrichment in the full accessible regions.

### RNA-seq sample preparation and data analysis

RNA was extracted using the RNeasy Mini Kit (#74136, Qiagen). Libraries were prepared using TruSeq Stranded Total RNA Library Prep Kit with Ribo-Zero Gold (Illumina) for experiments with PAEC exposed to LSS vs ST, caMEK5 vs GFP, and KLF2/4 RNAi studies, and using QuantSeq 3′ mRNA-Seq Library Prep Kit REV for Illumina (#016.96, Lexogen) for the DUSP5 RNAi stufy. Sequencing on the Illumina HiSeq 4000 yielded an average of approximately 30 million uniquely mapped reads for total RNA-Seq, and 6 million uniquely mapped reads per sample for mRNA-Seq for each (Stanford Center for Genomics and Personalized Medicine, supported by NIH grant S10OD020141). The resulting data were aligned to the human genome (GRCh37.p13) by STAR v.2.5.4b (https://github.com/alexdobin/STAR)^75^. The aligned transcripts were quantitated based on features in the GENCODE annotation database (GRCh37, version 19) by RSEM v. 1.3.1 (http://deweylab.biostat.wisc.edu/rsem/)^76^. Differentially expressed genes were detected using DESeq2 v. 1.20.0 (https://bioconductor.org/packages/release/bioc/html/DESeq2.html)^74^ with a P-value cutoff of <0.1. Functional enrichment for the differentially expressed genes was performed using Metascape^77^.

### ChIP-seq sample preparation and data analysis

For KLF4, H3K4me1 and H3K27ac ChIP-Seq, cells were trypsinized and cross-linked with 1% formaldehyde (EMD Millipore) for 10 min at RT. To quench the formaldehyde, 2 M glycine (ThermoFisher Scientific) was added and incubated for 5 min at room temperature. For BRG1 ChIP-Seq, cells were first cross-linked using 2 mM of DSG (Pierce) for 45 min at RT, washed with PBS, and then cross-linked with 1 % formaldehyde as described above. Cells were washed with ice cold PBS twice, snap-frozen and stored at −80°C. For ChIP-DNA preparation, cells were thawed by adding PBS and incubated at 4°C with rotation. Cells were treated with hypotonic buffer (20 mM HEPES pH 7.9, 10 mM KCl, 1 mM EDTA pH 8.0,10% glycerol) for 10 min on ice in the presence of protease inhibitors (G6521, Promega), then were homogenized using a glass homogenizer. Nuclear pellets were resuspended in RIPA buffer (Millipore) and incubated for 30 min on ice. Chromatin corresponding to 20 million cells for transcription factors, or 5 million cells for histone modifications was sheared with SFX250 Sonifier (Branson) and immunoprecipitated with antibodies targeting H3K27ac (#8173, Cell signaling Technology), H3K4me1 (#5326, Cell Signaling Technology), KLF4 (sc20691, Santa Cruz Biotechnology) and BRG1 (A303-877A, Bethyl Laboratories) at 4°C overnight on a nutator. For the input sample, 100 μl of sheared nuclear lysate was removed and stored overnight at 4°C. The next day, protein A/G agarose beads (Millipore) were added to the chromatin-antibody complex and incubated for one hour at 4°C on a nutator, after which the beads were eluted with SDS buffer (Santa Cruz Biotechnology) and incubated at 65°C for 10 min. Supernatant containing ChIP-DNA was reverse crosslinked by incubating overnight at 65°C. On the third day, ChIP-DNA was treated with RNase A (Qiagen) and proteinase K (ThermoFisher Scientific) and then purified. The ChIP-DNA samples were end repaired using End-It DNA End Repair Kit (Lucigen) and A-tailed using Klenow Fragment and dATP (New England Biolabs). Illumina TruSeq adapters (Illumina) were ligated using LigaFast (#M8221, Promega) and size-selected by gel extraction before PCR amplification. The purified libraries were subjected to 2×151 paired end sequencing on the Illumina HiSeq 4000 to obtain an average of approximately 26 million uniquely mapped reads for each sample (Stanford Center for Genomics and Personalized Medicine, supported by NIH grant S10OD020141). The resulting data were processed using the Kundaje Lab ChIP-seq processing pipeline (https://github.com/kundajelab/chipseq_pipeline). Briefly, this pipeline takes FASTQ files as input and outputs peak calls (bound regions; BR). Alignments to BR were counted using DiffBind 2.4.8 (https://rdrr.io/bioc/DiffBind/) to produce a count matrix. Differentially bound regions (DBR) were detected using DESeq2 (https://bioconductor.org/packages/release/bioc/html/DESeq2.html)^74^ with a P-value cutoff of <0.1. Data visualization was performed with IGV Genome Browser^78^.

### Immunofluorescence

PAEC were cultured on flow slides (μ-Slide I 0.4 mm ibiTreat; Ibidi), washed with PBS and fixed with ice-cold methanol at −20°C for 30 min. Methanol was aspirated and the slides were rehydrated with PBS at room temperature for 10 min. After washing with PBS, slides were blocked with 5% normal donkey serum and 2% BSA (Sigma Aldrich) in PBS at room temperature for 1 h. Incubation with primary antibodies targeting KLF4 (1:100, sc20691, Santa Cruz Biotechnology), ATF2 (1:100 sc-242, Santa Cruz Biotechnology) and ETS1 (1:100, sc55581, Santa Cruz Biotechnology) were carried out in the blocking buffer at 4°C overnight, and secondary antibody incubations (Alexa Fluor, ThermoFisher Scientific) in the blocking buffer at room temperature for 1 h. Slides were mounted with DAPI Fluoromount-G (DAPI, 4,6-diamidino-2-phenylindole) (SouthernBiotech). Stained slides were imaged using Leica Application Suite X software on a Leica Sp8 (Leica). Quantification of the nuclear fluorescence intensities was performed using ImageJ.

### Reverse Transcription (RT-) and ATAC-qPCR

For RT-qPCR, total RNA was extracted and purified using the Quick-RNA MiniPrep Kit (Zymo Research). The quantity and quality of RNA was determined using a spectrophotometer. RNA was reverse transcribed using the High Capacity RNA to cDNA Kit (Applied Biosystems) according to the manufacturer’s instructions. For ATAC-qPCR, cells were processed as described above for ATAC-Seq. For qPCR, nuclei were isolated from 50,000 cells and sequencing adapters were transposed for 30 min at 37°C using 2.5 μL of TDE1 (Nextera Tn5 transposase, Illumina). The reaction was terminated and purified using MinElute reaction cleanup (Qiagen) and used as template. In both cases, qPCR was performed using 1 μL of 5 μM Powerup SYBR green PCR Master Mix (Applied Biosystems), 2 μL of dH2O and 2 μL of cDNA sample in a 10 μL reaction. Each measurement was carried out in a duplicate using a CFX384 Real-Time System (Bio-Rad). The PCR conditions were: 95°C for 2 min, followed by 40 cycles of 95°C for 15 s, and 60°C for 60 s. Primer sequences used are listed in **Supplementary Table S2**. Gene expression levels were normalized to β-actin, and accessibility changes were normalized to GAPDH.

### Affinity Purification followed by Mass Spectrometry (AP-MS)

Nuclear fractionation and AP were performed using the Nuclear Complex Co-IP kit (ActiveMotif) according to manufacturer’s instructions. Briefly, cells were trypsinized and washed with cold PBS. The cell pellets were resuspended in hypotonic buffer and incubated on ice for 15 min. Detergent was added and the suspension was centrifuged at for 30 s at 14,000 g in a pre-cooled centrifuge. The nuclear pellet was resuspended in complete digestion buffer with 0.75 μL of enzymatic shearing cocktail and incubated at 37°C for 10 min. 0.5M EDTA (3 μL) was added to stop the reaction. The suspension was placed on ice for 5 min and then centrifuged for 10 min at 14,000 g. For AP, the supernatant was pre-cleared by adding 30 μL of Dynabeads Protein G (10004D, ThermoFisher Scientific) and incubated on a rotator at 4°C for 1 h. The beads were removed and the supernatant was used for subsequent AP. Antibodies (2 μg) targeting either KLF (sc166238, Santa Cruz Biotechnology) or FLAG (F7425, Millipore Sigma) were added per 1 mg of pre-cleared supernatant and incubated on a rotator at 4°C overnight. The next day, samples were incubated with 30 μL of Dynabeads Protein G (10004D, ThermoFisher Scientific) on a rotator at 4°C for 1h. After three successive washes with ice-cold washing buffer, the proteins were eluted from the beads using 100 μL of IgG Elution Buffer (ThermoFisher Scientific) at a gentle vortex at room temperature (RT) for 7 min. The eluate was then immediately neutralized with 1:10 1M Tris-HCl, pH 8.5. For MS analysis, samples were reduced with 5 mM DTT in 120 μL of 50 mM ammonium bicarbonate. Following reduction, proteins were alkylated using 10 mM acrylamide for 30 min at room temperature to cap cysteines. Digestion was performed using Trypsin/LysC (Promega) overnight at 37°C. Following digestion and acid quenching, samples were passed over HILIC resin (Resyn Biosciences), dried in a speed vac and then reconstituted in 10 μL reconstitution buffer (2% acetonitrile with 0.1% Formic acid); 3 μL of the reconstituted peptides were injected on the instrument.

All mass spectrometry experiments were performed using an Orbitrap Fusion Tribrid mass spectrometer (ThermoFisher Scientific) with an attach Acquity M-Class UPLC (Waters Corporation) liquid chromatograph. A pulled-and-packed fused silica C18 reverse phase column containing Dr. Maisch 1.8 μm C18 beads and a length of ∼25 cm was used over a 80 min gradient. A flow rate of 300 nL/min was used with the mobile phase A consisting of aqueous 0.2% formic acid and mobile phase B consisting of 0.2% formic acid in acetonitrile. Peptides were directly injected onto the analytical column. The mass spectrometer was operated in a data dependent fashion, with MS1 survey spectra collected in the Orbitrap and MS2 fragmentation using CID for in the ion trap.

For data analysis, the .raw data files were processed using Byonic v2.14.27 (Protein Metrics) to identify peptides and infer proteins against the human UniProt database containing isoforms concatenated with synthesized sequences. Proteolysis was assumed to be tryptic in nature and allowed for ragged n-terminal digestion and up to two missed cleavage sites. Precursor mass accuracies were held within 12 ppm, with MS/MS fragments held to a 0.4 Da mass accuracy. Proteins were held to a false discovery rate of 1%, using standard approaches^79^.

### Proximity ligation assays (PLA)

PLA in cultured PAEC were performed using Duolink PLA protein detection technology (Millipore Sigma) according to manufacturer’s instructions. Briefly, slides with PAEC were crosslinked with 4% PFA for 10 min, washed and incubated with Duolink Blocking solution for 1 h at 37°C. Slides were incubated with rabbit antibodies targeting KLF4 (1:100, sc20691, Santa Cruz Biotechnology) and mouse antibodies targeting BRG1 (1:75, sc17796, Santa Cruz Biotechnology) or SMARCC2 (1:75, sc17838, Santa Cruz Biotechnology) overnight at 4°C. Rabbit and mouse IgG were used as controls. The next day, slides were washed, and then incubated with the Duolink PLUS and MINUS probes for 1 h at 37°C. After washing the slides, the probes were ligated for 30 min at 37°C. Slides were washed and rolling circle amplification was performed for 100 min at 37°C, after which slides underwent final washes and were mounted using Duolink in situ mounting medium with DAPI (Millipore Sigma). Stained slides were imaged using Leica Application Suite X software on a Leica Sp8 (Leica). Quantification of the average number of foci per nucleus was performed using ImageJ.

For PLA in rat lung tissue, the protocol was modified using longer incubation times to allow reagents to fully penetrate the tissues. In brief, lungs from healthy rats were flushed and perfusion fixed with 4% PFA for 30 min on ice, washed with PBS and dehydrated in methanol. Following rehydration, tissue was cut in 350 μM thick sections. Sections were bleached using 3% H2O2 for 1 h at RT, washed with 0.5% triton-PBS and blocked using 5% DS 5% BSA in 0.5% triton-PBS for 3 h at RT. Sections were incubated with primary antibodies targeting KLF4 (1:50, sc20691, Santa Cruz Biotechnology) and BRG1 (1:30, sc17796, Santa Cruz Biotechnology) in 3% DS 1% BSA in 0.1% triton-PBS overnight at 4°C. The following day, sections were washed, blocked using 3% DS 1% BSA in 0.1% triton-PBS (dilution buffer) for 4 h at RT, after which they were incubated with the Duolink PLUS and MINUS probes overnight at 4°C. On the next day, sections were washed, and the probes were ligated for 2 h at 37°C. After washing, rolling circle amplification was performed for 3 h at 37°C, after which slides underwent final washes. Sections were post-fixed using 4% PFA for 25 min at RT, washed and blocked with anti-Rabbit IgG in dilution buffer at RT. Sections were then incubated with FITC-conjugated anti-aSMA antibodies (1:400, F377, Sigma-Aldrich) and antibodies targeting vWF (1:500, ab6994, Abcam) overnight at 4°C. After washing, sections were incubated with anti-Rabbit Alexa Fluor 647-conjugated secondary antibodies (1:200, A32795, Thermo Fisher Scientific) for 3 h at RT, washed in triton-PBS with DAPI for 2 h at RT, and post-fixed with 4% PFA for 30 min at RT. Finally, sections were washed, dehydrated in methanol, and cleared using Benzyl Alcohol/ Benzyl Benzoate (BABB). Sections were imaged using Leica Application Suite X software on a Leica Sp8 (Leica). Three-dimensional reconstructions were made with Imaris version 9.3.0 (Bitplane).

### H3K27ac HiC Chromatin Immunoprecipitation (HiChIP)

H3K27ac HiChIP was performed as previously described^19^. PAEC were crosslinked in 1% formaldehyde for 10 min at room temperature and then quenched by 125 mM Glycine for 5 min at RT. Nuclei were isolated from 1 million crosslinked cells by 30 min of lysis at 4°C. Nuclei were permeabilized in 0.5% SDS for 10 min at 62°C and quenched using Triton X-100 for 15 min at 37°C. MboI restriction enzyme (R0147, New England Biolabs) was added to digest chromatin for 2 h at 37°C and then heat-inactivated for 20 min at 62°C. Klenow was then used to fill in restriction fragment overhangs and mark the DNA ends with biotin (M0210, New England Biolabs). Proximity ligation contact (PLC) pellets were then created by incubation with DNA ligase for 4 h at room temperature followed by centrifugation. PLC pellets were then sonicated and immunoprecipitated using H3K27ac antibody (#8173, Cell signaling Technology) as previously described. The eluted fragments labeled by biotin were then captured by streptavidin bead pull-down. DNA was then adaptor-labeled using Tn5 transposase (Illumina) and subjected to PCR amplification. Samples were then sequenced by 2×101 paired-end sequencing on an Illumina NovaSeq 6000 to an average yield of 200 million reads per sample. The resulting data were filtered for duplicate reads, aligned to the hg19 genome, and filtered for valid interactions using the HiC-Pro pipeline v.2.11.1 (https://github.com/nservant/HiC-Pro) using the default settings. FitHiChIP (https://github.com/ay-lab/FitHiChIP) was then used to determine statistically significant interactions using default settings, with the exception of allowing interactions with a minimum size of 1Kb. The diffLoop package (https://github.com/aryeelab/diffloop) was used to test for differential interactions between conditions, and to infer gene enhancer to promoter relationships.

### Activity-by-Contact Model (ABC)

The ABC v0.2 pipeline was cloned from the GitHub repository (https://github.com/broadinstitute/ABC-Enhancer-Gene-Prediction/). First, ATAC peaks were called by MACS2 v.2.1.2 (https://github.com/taoliu/MACS) from each ATAC BAM file with a P-value cutoff of 0.1. Candidate enhancer regions were then defined by the ABC script makeCandidateRegions.py, which: (1) resized each peak to be 250 base pairs centered on the peak summit. (2) Counted ATAC-seq reads in each peak and retained the top 150,000 peaks with the most read counts. (3) removed any regions that are blacklisted due to known propensity for errors (hg19-blacklist.v2.bed from https://github.com/Boyle-Lab/Blacklist) and (4) merged any overlapping regions. Enhancer activity was then quantified by the ABC script run.neighborhoods.py, which counted ATAC-seq and H3K27ac ChIP-seq reads in the candidate enhancer regions that were generated in the previous step, gene bodies, and promoter regions. Lastly, the ABC score was calculated using the ABC script predict.py; which combined information from the enhancer and promoter activities, calculated in the previous step, with contact frequency data from average Hi-C profiles of 10 cell lines (ftp://ftp.broadinstitute.org/outgoing/lincRNA/average_hic/average_hic.v2.191020.tar.gz. The default threshold of 0.02 was applied which corresponds to approximately 70% recall and 60% precision^21^.

### Data and Software Availability

ATAC-Seq, RNA-Seq, ChIP-seq and HiChIP data is deposited in the Gene Expression Omnibus (GEO) under accession number GSE152900. AP-MS data will be deposited in ProteomeXchange. The datasets generated and/or analyzed during the current study are available from the corresponding author on reasonable request.

### Statistical analysis

Values from multiple experiments are shown as arithmetical mean ± SEM. Statistical significance was determined using unpaired two-tailed Student’s *t* test. Correlations were calculated by Pearson R test. A *P*-value of <0.05 was considered significant. The number of samples in each group, the statistical test used and the statistical significance is indicated in the figures or figure legends. Data were analyzed using Prism version 8.4 (Graphpad).

## References

1. Friedman, M.H., Bargeron, C.B., Deters, O.J., Hutchins, G.M. & Mark, F.F. Correlation between wall shear and intimal thickness at a coronary artery branch. Atherosclerosis 68, 27–33 (1987).

2. Ku, D.N., Giddens, D.P., Zarins, C.K. & Glagov, S. Pulsatile flow and atherosclerosis in the human carotid bifurcation. Positive correlation between plaque location and low oscillating shear stress. Arteriosclerosis 5, 293–302 (1985).

3. Cybulsky, M.I. & Gimbrone, M.A., Jr. Endothelial expression of a mononuclear leukocyte adhesion molecule during atherogenesis. Science 251, 788–791 (1991).

4. Davies, P.F., Civelek, M., Fang, Y. & Fleming, I. The atherosusceptible endothelium: endothelial phenotypes in complex haemodynamic shear stress regions in vivo. Cardiovasc Res 99, 315–327 (2013).

5. Malek, A.M., Alper, S.L. & Izumo, S. Hemodynamic shear stress and its role in atherosclerosis. Jama 282, 2035–2042 (1999).

6. Tzima, E. et al. A mechanosensory complex that mediates the endothelial cell response to fluid shear stress. Nature 437, 426–431 (2005).

7. Hahn, C. & Schwartz, M.A. Mechanotransduction in vascular physiology and atherogenesis. Nat Rev Mol Cell Biol 10, 53–62 (2009).

8. Davies, P.F. Flow-mediated endothelial mechanotransduction. Physiol Rev 75, 519–560 (1995).

9. Dekker, R.J. et al. Prolonged fluid shear stress induces a distinct set of endothelial cell genes, most specifically lung Kruppel-like factor (KLF2). Blood 100, 1689–1698 (2002).

10. Parmar, K.M. et al. Integration of flow-dependent endothelial phenotypes by Kruppel-like factor 2. J Clin Invest 116, 49–58 (2006).

11. Hamik, A. et al. Kruppel-like factor 4 regulates endothelial inflammation. J Biol Chem 282, 13769–13779 (2007).

12. Villarreal, G., Jr. et al. Defining the regulation of KLF4 expression and its downstream transcriptional targets in vascular endothelial cells. Biochem Biophys Res Commun 391, 984–989 (2010).

13. Sangwung, P. et al. KLF2 and KLF4 control endothelial identity and vascular integrity. JCI Insight 2, e91700 (2017).

14. Pietra, G.G. et al. Histopathology of primary pulmonary hypertension. A qualitative and quantitative study of pulmonary blood vessels from 58 patients in the National Heart, Lung, and Blood Institute, Primary Pulmonary Hypertension Registry. Circulation 80, 1198–1206 (1989).

15. Rabinovitch, M. Molecular pathogenesis of pulmonary arterial hypertension. J Clin Invest 122, 4306–4313 (2012).

16. Deng, Z. et al. Familial primary pulmonary hypertension (gene PPH1) is caused by mutations in the bone morphogenetic protein receptor-II gene. Am J Hum Genet 67, 737–744 (2000).

17. Lane, K.B. et al. Heterozygous germline mutations in BMPR2, encoding a TGF-beta receptor, cause familial primary pulmonary hypertension. Nat Genet 26, 81–84 (2000).

18. Gupta, R.M. et al. A Genetic Variant Associated with Five Vascular Diseases Is a Distal Regulator of Endothelin-1 Gene Expression. Cell 170, 522–533.e515 (2017).

19. Mumbach, M.R. et al. HiChIP: efficient and sensitive analysis of protein-directed genome architecture. Nat Methods 13, 919–922 (2016).

20. Mumbach, M.R. et al. Enhancer connectome in primary human cells identifies target genes of disease-associated DNA elements. Nat Genet 49, 1602–1612 (2017).

21. Fulco, C.P. et al. Activity-by-contact model of enhancer-promoter regulation from thousands of CRISPR perturbations. Nat Genet 51, 1664–1669 (2019).

22. Tang, B.T. et al. Wall shear stress is decreased in the pulmonary arteries of patients with pulmonary arterial hypertension: An image-based, computational fluid dynamics study. Pulm Circ 2, 470–476 (2012).

23. McLaughlin, F. et al. Combined genomic and antisense analysis reveals that the transcription factor Erg is implicated in endothelial cell differentiation. Blood 98, 3332–3339 (2001).

24. De Val, S. et al. Combinatorial regulation of endothelial gene expression by ets and forkhead transcription factors. Cell 135, 1053–1064 (2008).

25. Ohnesorge, N. et al. Erk5 activation elicits a vasoprotective endothelial phenotype via induction of Kruppel-like factor 4 (KLF4). J Biol Chem 285, 26199–26210 (2010).

26. Moonen, J.R. et al. Endothelial-to-mesenchymal transition contributes to fibro-proliferative vascular disease and is modulated by fluid shear stress. Cardiovasc Res 108, 377–386 (2015).

27. Mandl, M., Slack, D.N. & Keyse, S.M. Specific inactivation and nuclear anchoring of extracellular signal-regulated kinase 2 by the inducible dual-specificity protein phosphatase DUSP5. Mol Cell Biol 25, 1830–1845 (2005).

28. Bhattacharyya, S., Chandra, V., Vijayanand, P. & Ay, F. Identification of significant chromatin contacts from HiChIP data by FitHiChIP. Nat Commun 10, 4221 (2019).

29. Giaid, A. et al. Expression of endothelin-1 in the lungs of patients with pulmonary hypertension. N Engl J Med 328, 1732–1739 (1993).

30. Channick, R.N. et al. Effects of the dual endothelin-receptor antagonist bosentan in patients with pulmonary hypertension: a randomised placebo-controlled study. Lancet 358, 1119–1123 (2001).

31. Salmon, M., Gomez, D., Greene, E., Shankman, L. & Owens, G.K. Cooperative binding of KLF4, pELK-1, and HDAC2 to a G/C repressor element in the SM22alpha promoter mediates transcriptional silencing during SMC phenotypic switching in vivo. Circ Res 111, 685–696 (2012).

32. Illi, B. et al. Shear stress-mediated chromatin remodeling provides molecular basis for flow-dependent regulation of gene expression. Circ Res 93, 155–161 (2003).

33. Chen, W., Bacanamwo, M. & Harrison, D.G. Activation of p300 histone acetyltransferase activity is an early endothelial response to laminar shear stress and is essential for stimulation of endothelial nitric-oxide synthase mRNA transcription. J Biol Chem 283, 16293–16298 (2008).

34. Chen, Z. et al. Shear stress, SIRT1, and vascular homeostasis. Proc Natl Acad Sci U S A 107, 10268–10273 (2010).

35. Zampetaki, A. et al. Histone deacetylase 3 is critical in endothelial survival and atherosclerosis development in response to disturbed flow. Circulation 121, 132–142 (2010).

36. Lee, D.Y. et al. Role of histone deacetylases in transcription factor regulation and cell cycle modulation in endothelial cells in response to disturbed flow. Proc Natl Acad Sci U S A 109, 1967–1972 (2012).

37. Maleszewska, M., Vanchin, B., Harmsen, M.C. & Krenning, G. The decrease in histone methyltransferase EZH2 in response to fluid shear stress alters endothelial gene expression and promotes quiescence. Angiogenesis 19, 9–24 (2016).

38. Dunn, J. et al. Flow-dependent epigenetic DNA methylation regulates endothelial gene expression and atherosclerosis. J Clin Invest 124, 3187–3199 (2014).

39. Zhou, J., Li, Y.S., Wang, K.C. & Chien, S. Epigenetic Mechanism in Regulation of Endothelial Function by Disturbed Flow: Induction of DNA Hypermethylation by DNMT1. Cell Mol Bioeng 7, 218–224 (2014).

40. Jiang, Y.Z., Manduchi, E., Stoeckert, C.J., Jr. & Davies, P.F. Arterial endothelial methylome: differential DNA methylation in athero-susceptible disturbed flow regions in vivo. BMC Genomics 16, 506 (2015).

41. Jiang, Y.Z. et al. Hemodynamic disturbed flow induces differential DNA methylation of endothelial Kruppel-Like Factor 4 promoter in vitro and in vivo. Circ Res 115, 32–43 (2014).

42. He, M. et al. Atheroprotective Flow Upregulates ITPR3 (Inositol 1,4,5-Trisphosphate Receptor 3) in Vascular Endothelium via KLF4 (Kruppel-Like Factor 4)-Mediated Histone Modifications. Arterioscler Thromb Vasc Biol 39, 902–914 (2019).

43. Kumar, S., Kim, C.W., Simmons, R.D. & Jo, H. Role of flow-sensitive microRNAs in endothelial dysfunction and atherosclerosis: mechanosensitive athero-miRs. Arterioscler Thromb Vasc Biol 34, 2206–2216 (2014).

44. Wu, W. et al. Flow-Dependent Regulation of Kruppel-Like Factor 2 Is Mediated by MicroRNA-92a. Circulation 124, 633–641 (2011).

45. Fang, Y. & Davies, P.F. Site-specific microRNA-92a regulation of Kruppel-like factors 4 and 2 in atherosusceptible endothelium. Arterioscler Thromb Vasc Biol 32, 979–987 (2012).

46. Man, H.S.J. et al. Angiogenic patterning by STEEL, an endothelial-enriched long non-coding RNA. Proc Natl Acad Sci U S A 115, 2401–2406 (2018).

47. Miao, Y. et al. Enhancer-associated long non-coding RNA LEENE regulates endothelial nitric oxide synthase and endothelial function. Nat Commun 9, 292 (2018).

48. Orphanides, G., Lagrange, T. & Reinberg, D. The general transcription factors of RNA polymerase II. Genes Dev 10, 2657–2683 (1996).

49. Banerji, J., Rusconi, S. & Schaffner, W. Expression of a beta-globin gene is enhanced by remote SV40 DNA sequences. Cell 27, 299–308 (1981).

50. Osterwalder, M. et al. Enhancer redundancy provides phenotypic robustness in mammalian development. Nature 554, 239–243 (2018).

51. Reyes-Palomares, A. et al. Remodeling of active endothelial enhancers is associated with aberrant gene-regulatory networks in pulmonary arterial hypertension. Nat Commun 11, 1673 (2020).

52. Takahashi, K. & Yamanaka, S. Induction of pluripotent stem cells from mouse embryonic and adult fibroblast cultures by defined factors. Cell 126, 663–676 (2006).

53. Iwafuchi-Doi, M. & Zaret, K.S. Pioneer transcription factors in cell reprogramming. Genes Dev 28, 2679–2692 (2014).

54. Soufi, A. et al. Pioneer transcription factors target partial DNA motifs on nucleosomes to initiate reprogramming. Cell 161, 555–568 (2015).

55. Soufi, A., Donahue, G. & Zaret, K.S. Facilitators and impediments of the pluripotency reprogramming factors’ initial engagement with the genome. Cell 151, 994–1004 (2012).

56. Kwon, H., Imbalzano, A.N., Khavari, P.A., Kingston, R.E. & Green, M.R. Nucleosome disruption and enhancement of activator binding by a human SW1/SNF complex. Nature 370, 477–481 (1994).

57. Kingston, R.E. & Narlikar, G.J. ATP-dependent remodeling and acetylation as regulators of chromatin fluidity. Genes Dev 13, 2339–2352 (1999).

58. Burns, L.G. & Peterson, C.L. The yeast SWI-SNF complex facilitates binding of a transcriptional activator to nucleosomal sites in vivo. Mol Cell Biol 17, 4811–4819 (1997).

59. Cosma, M.P., Tanaka, T. & Nasmyth, K. Ordered recruitment of transcription and chromatin remodeling factors to a cell cycle- and developmentally regulated promoter. Cell 97, 299–311 (1999).

60. Ryan, M.P., Jones, R. & Morse, R.H. SWI-SNF complex participation in transcriptional activation at a step subsequent to activator binding. Mol Cell Biol 18, 1774–1782 (1998).

61. Li, Z. et al. Foxa2 and H2A.Z mediate nucleosome depletion during embryonic stem cell differentiation. Cell 151, 1608–1616 (2012).

62. Khavari, P.A., Peterson, C.L., Tamkun, J.W., Mendel, D.B. & Crabtree, G.R. BRG1 contains a conserved domain of the SWI2/SNF2 family necessary for normal mitotic growth and transcription. Nature 366, 170–174 (1993).

63. Chronis, C. et al. Cooperative Binding of Transcription Factors Orchestrates Reprogramming. Cell 168, 442–459.e420 (2017).

64. Alver, B.H. et al. The SWI/SNF chromatin remodelling complex is required for maintenance of lineage specific enhancers. Nat Commun 8, 14648 (2017).

65. Kidder, B.L., Palmer, S. & Knott, J.G. SWI/SNF-Brg1 regulates self-renewal and occupies core pluripotency-related genes in embryonic stem cells. Stem Cells 27, 317–328 (2009).

66. Mak, A.B. et al. A lentiviral functional proteomics approach identifies chromatin remodeling complexes important for the induction of pluripotency. Mol Cell Proteomics 9, 811–823 (2010).

67. Fledderus, J.O. et al. Prolonged shear stress and KLF2 suppress constitutive proinflammatory transcription through inhibition of ATF2. Blood 109, 4249–4257 (2007).

68. Dekker, J., Rippe, K., Dekker, M. & Kleckner, N. Capturing chromosome conformation. Science 295, 1306–1311 (2002).

69. Lieberman-Aiden, E. et al. Comprehensive mapping of long-range interactions reveals folding principles of the human genome. Science 326, 289–293 (2009).

70. Rao, S.S. et al. A 3D map of the human genome at kilobase resolution reveals principles of chromatin looping. Cell 159, 1665–1680 (2014).

71. Di Giammartino, D.C. et al. KLF4 is involved in the organization and regulation of pluripotency-associated three-dimensional enhancer networks. Nat Cell Biol 21, 1179–1190 (2019).

72. Maurano, M.T. et al. Systematic localization of common disease-associated variation in regulatory DNA. Science 337, 1190–1195 (2012).

73. Buenrostro, J.D., Giresi, P.G., Zaba, L.C., Chang, H.Y. & Greenleaf, W.J. Transposition of native chromatin for fast and sensitive epigenomic profiling of open chromatin, DNA-binding proteins and nucleosome position. Nat Methods 10, 1213–1218 (2013).

74. Love, M.I., Huber, W. & Anders, S. Moderated estimation of fold change and dispersion for RNA-seq data with DESeq2. Genome Biol 15, 550 (2014).

75. Dobin, A. et al. STAR: ultrafast universal RNA-seq aligner. Bioinformatics 29, 15–21 (2013).

76. Li, B. & Dewey, C.N. RSEM: accurate transcript quantification from RNA-Seq data with or without a reference genome. BMC Bioinformatics 12, 323 (2011).

77. Zhou, Y. et al. Metascape provides a biologist-oriented resource for the analysis of systems-level datasets. Nat Commun 10, 1523 (2019).

78. Thorvaldsdottir, H., Robinson, J.T. & Mesirov, J.P. Integrative Genomics Viewer (IGV): high-performance genomics data visualization and exploration. Brief Bioinform 14, 178–192 (2013).

79. Elias, J.E. & Gygi, S.P. Target-decoy search strategy for increased confidence in large-scale protein identifications by mass spectrometry. Nat Methods 4, 207–214 (2007).

