## Supplementary Data for "KLF4 Recruits SWI/SNF to Increase Chromatin Accessibility and Reprogram the Endothelial Enhancer Landscape under Laminar Shear Stress"

### Fig S1: KLF2 and KLF4 are induced in PAEC exposed to LSS.

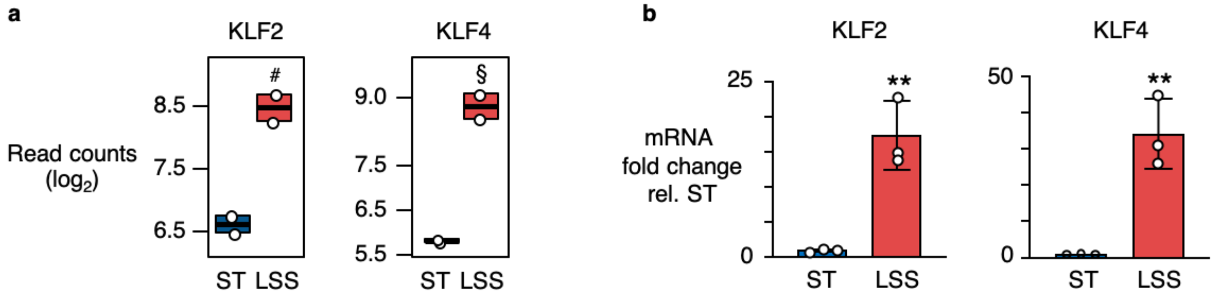


**a**, Box plot depicting the normalized read counts for *KLF2* and *KLF4* obtained by RNA-Seq from PAEC exposed to 15 dyn/cm$^2$ of LSS for 24h vs ST. *n*=2 experimental replicates. *P* values were determined by the Wald test with Benjamini–Hochberg adjustment. # *P*=5.28E-13, § *P*=8.00E-32. **b**, Bar graphs showing confirmation by RT-qPCR of the induction of *KLF2* and *KLF4* upon exposure to 15 dyn/cm$^2$ of LSS for 24h, normalized to ST control levels. *n*=3 experimental replicates. Bars represent mean ± s.e.m. ***P*<0.01 by Student's two-tailed *t*-test.

**Fig S2: Validation of KLF gain-and-loss of function and DUSP5 target genes.**

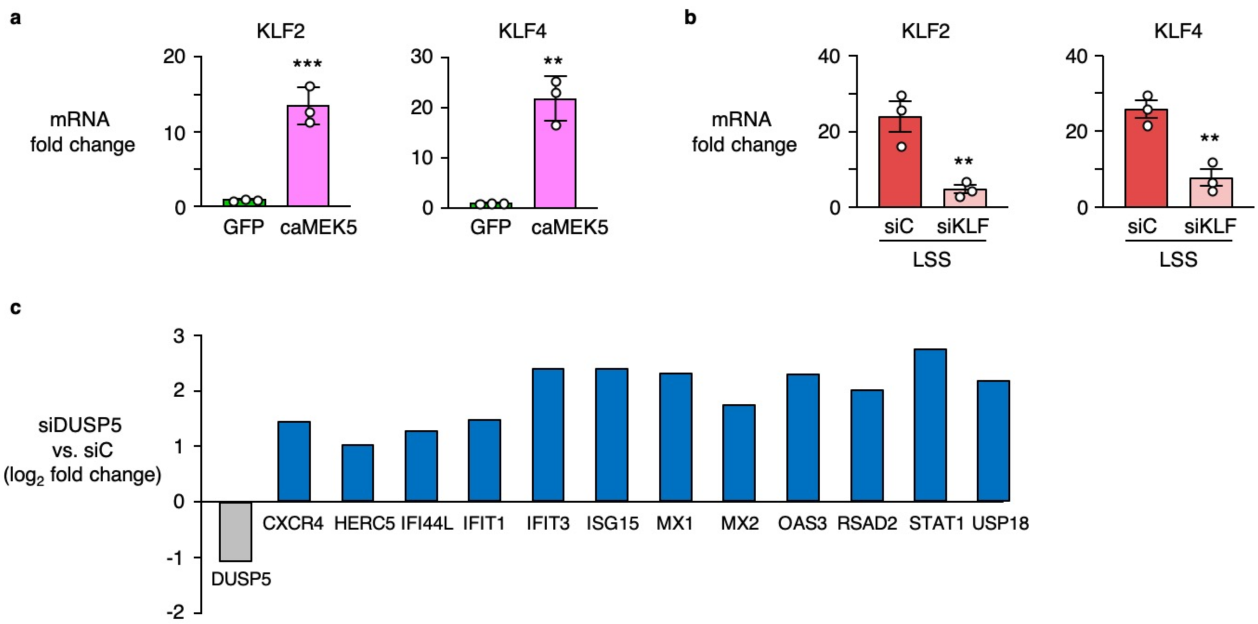


**a**, Bar graphs showing the expression of *KLF2* and *KLF4* by RT-qPCR in PAEC transduced with adenoviral vectors encoding caMEK5, normalized to GFP controls. $n$=3 experimental replicates. Bars represent mean ± s.e.m. **$P$<0.01, ***$P$<0.001, by Student's two-tailed $t$-test. **b**, Bar graphs showing the expression of *KLF2* and *KLF4* by RT-qPCR in PAEC under LSS conditions, treated with siRNA targeting *KLF2* and *KLF4* (siKLF) vs non-targeting negative controls (siC), normalized to Static siC. $n$=3 experimental replicates. Bars represent mean ± s.e.m. ** $P$< 0.01, by Student's two-tailed $t$-test. **c**, Bar graphs from RNA-Seq data in PAEC treated with siRNA targeting *DUSP5* prior to exposure to LSS, indicating de-repression of genes related to an antiviral response and activation of the interferon signaling pathway. $n$=2 experimental replicates. $P$ values were determined by the Wald test with Benjamini–Hochberg adjustment, all genes listed were significant using a 10% FDR threshold.

**Fig S3: Deep tissue imaging of rat lung tissues and validation of BRG1 loss-of-function.**

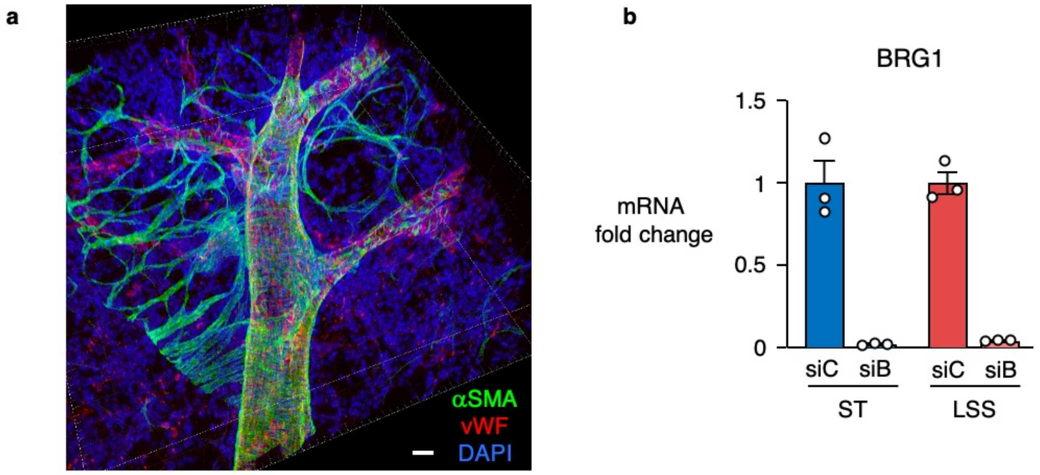


**a**, Representative image of a 3D reconstruction of a rat pulmonary artery. αSMA (green); vWF (red, pseudo-colour). Nuclei were stained with DAPI, (blue). Scale bar, 30 μm. **b**, Bar graphs showing the expression of BRG1 by RT-qPCR in PAEC treated with siRNA targeting BRG1 (siB) vs non-targeting negative controls (siC) under ST or LSS conditions, normalized to Static siC. *n*=3 experimental replicates. Bars represent mean ± s.e.m. ** P< 0.01, by Student's two-tailed *t*-test.

**Fig S4: KLF4 and BRG1 co-occupancy at AR, correlation between BRG1 occupancy and accessibility changes, and enrichment of BRG1 at KLF4 bound sites.**

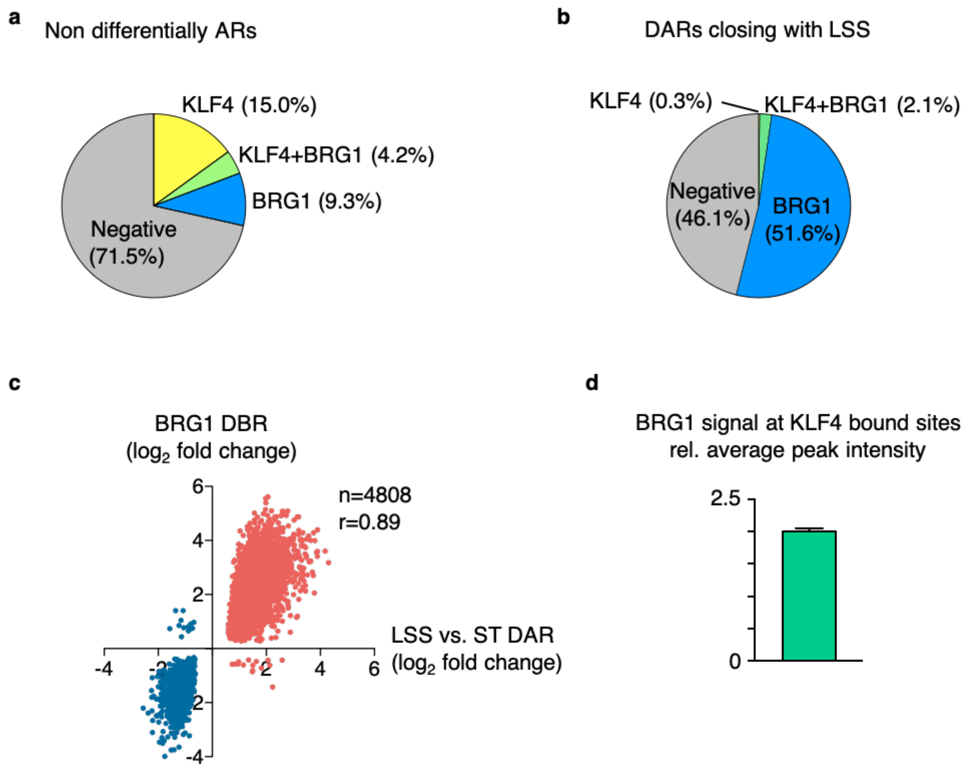


**a**, Pie chart showing the percentage of regions that are accessible under both LSS and ST, that are differentially enriched for KLF4 and/or BRG1. **b,** Pie chart showing enrichment for KLF4 and/or BRG1 in regions that have an increased accessibility under ST. **c,** Scatterplot showing the correlation between differential BRG1 binding (DBR) and accessibility changes at LSS vs ST DAR. r=0.89 with $P < 0.0001$ calculated using Pearson R test. **d,** Bar graph showing increased BRG1 levels at KLF4 co-occupied sites, relative to the average BRG1 intensity across all BRG1 peaks. *n*=2 experimental replicates.

**Table S1: Demographic and clinical data from donor control PAEC.**

| ID | Age (yr) | Gender | Race | Ethnicity | Cause of Death |
|---|---|---|---|---|---|
| PAEC-01 PROMO | 28 | Male | White | Unknown | Polytrauma |
| PAEC-02 PROMO | 19 | Male | White | Unknown | Cardiac arrythmia |
| PAEC-03 | 12 | Male | Unknown | Non-Hispanic | Head trauma from motor vehicle accident |
| PAEC-04 | 1 | Male | White | Non-Hispanic | Anoxia/drowning |
| PAEC-05 | 46 | Male | Asian | Unknown | Intracranial hemorrhage |
| PAEC-06 | 25 | Male | White | Non-Hispanic | Intracranial hemorrhage |
| PAEC-07 | 16 | Male | White | Non-Hispanic | Gunshot wound resulting in subarachnoid hemorrhage |

Primary human pulmonary artery endothelial cells (PAEC) were either commercially obtained (PAEC-01 PROMO and PAEC-02 PROMO; PromoCell), or harvested from unused donor control lungs obtained though the Pulmonary Hypertension Breakthrough Initiative (PHBI) funded by NIH (R24 HL123767) and the Cardiovascular Medical Research and Education Fund (CMREF; UL 1RR024986). De-identified demographic and clinical data were obtained from the PHBI Data Coordinating Center at the University of Michigan.

**Table S2: qPCR primers used for RT-qPCR and ATAC-qPCR.**

RT-qPCR primers

| Gene | Forward | Reverse |
|---|---|---|
| BMPR2 | CTGCGGCTGCTTCGCAGAAT | TGGTGTTGTGTCAGGAGGTGG |
| DUSP5 | ACAAATGGATCCCTGTGGAA | CCTCCCTTTTCCCTGACAC |
| EDN1 | ACGGAACAACGTGCTCGGGA | AGTGGGTTTCTCCCCGCCGT |
| HES2 | CGCATCAACCAGAGCCTGA | GAGCAGTTGGAGTTCTCCCG |
| KLF2 | CATCTGAAGGCGCATCTG | CGTGTGCTTTCGGTAGTGG |
| KLF4 | GGGAGAAGACACTGCGTCA | GGAAGCACTGGGGGAAGT |
| SMAD5 | TTGCTCAGCTTCTGGCTCAA | CCGGTGATATTCTGCTCCCC |

ATAC-qPCR primers

| Gene | Forward | Reverse |
|---|---|---|
| DUSP5 | CCTCTGCTTTAAATGCCCGG | TTCTTGCCCCTGTAACCACC |
| GAPDH | CATCTCAGTCGTTCCCAAAGT | TTCCCAGGACTGGACTGT |
